# Pro-inflammatory mediators sensitise Transient Receptor Potential Melastatin 3 cation channel (TRPM3) signalling in mouse sensory neurons

**DOI:** 10.1101/2024.09.11.612393

**Authors:** Javier Aguilera-Lizarraga, Tony K. Lim, Luke A. Pattison, Luke W. Paine, David C. Bulmer, Ewan St. John Smith

## Abstract

Pro-inflammatory mediators can directly activate pain-sensing neurons, known as nociceptors. Additionally, these mediators can potentiate or sensitise ion channels and receptors expressed by these cells through transcriptional and post-translational modulation, leading to nociceptor hypersensitivity. A well-characterised group of ion channels that subserve nociceptor sensitisation is the transient receptor potential (TRP) superfamily of cation channels. For example, the roles of TRP channels vanilloid 1 (TRPV1) and ankyrin 1 (TRPA1) in nociceptor sensitisation and inflammatory pain have been extensively documented. In the case of TRP melastatin 3 (TRPM3), however, despite the increasing recognition of this channel’s role in inflammatory pain, the mechanisms driving its sensitisation during inflammation remain poorly understood. Here, we found that an inflammatory soup of bradykinin, interleukin 1β (IL-1β) and tumour necrosis factor α (TNFα) sensitised TRPM3 function in isolated mouse sensory neurons; IL-1β and TNFα, but not bradykinin, independently potentiated TRPM3 function. TRPM3 expression and translocation to the membrane remained unchanged upon individual or combined exposure to these inflammatory mediators, which suggests post-translational modification occurs. Finally, using the model of complete Freund’s adjuvant-induced knee inflammation, we found that pharmacological blockade of TRPM3 does not alleviate inflammatory pain, which contrasts with previous reports using different pain models. We propose that the nuances of the immune response may determine the relative contribution of TRPM3 to nociceptive signalling in different neuro-immune contexts. Collectively, our findings improve insight into the role of TRPM3 sensitisation in inflammatory pain.

Graphical abstract

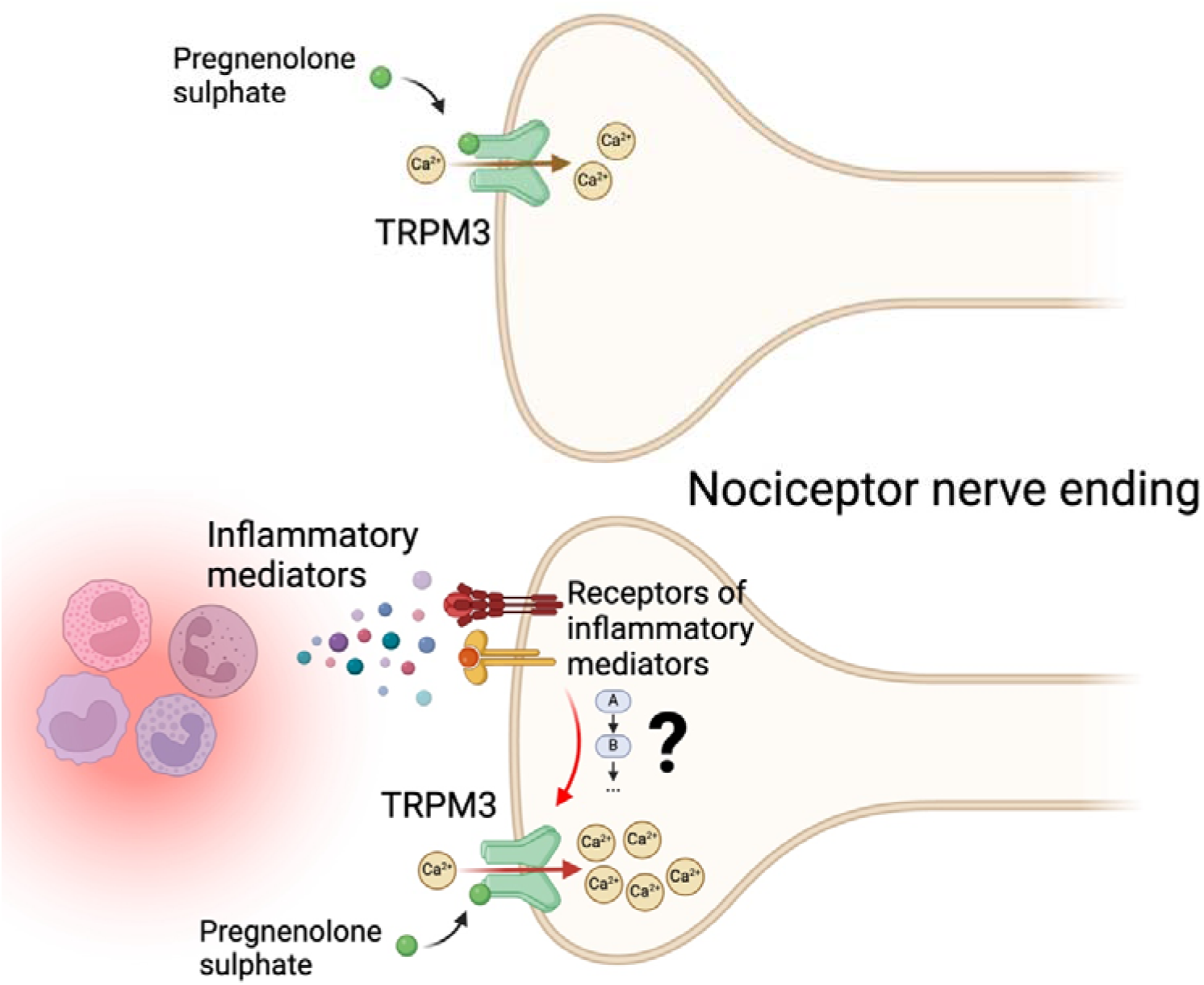

## 1. Introduction

Activation of the nervous system is a crucial and significant consequence of inflammation. Indeed, one of the primary effects of inflammatory states is the development of pain (Pinho-Ribeiro et al., 2017), which is due to activation of nociceptors, specialised ‘pain-sensing’ neurons. Nociceptors have their cell bodies in the dorsal root ganglia (DRG) and express specific receptors that recognise inflammatory cytokines and other immune-derived mediators (Cook et al., 2018). During pathological inflammatory states, pro-inflammatory mediators can disrupt normal nociceptor function, causing them to become hypersensitive, a process known as peripheral sensitisation (Hucho and Levine, 2007; Pethö and Reeh, 2012). Nociceptor hyperexcitability underpins allodynia, hyperalgesia, and/or spontaneous pain, and may ultimately lead to the development of chronic pain (Gold and Gebhart, 2010). Despite many advances in the field, the plethora of molecular mechanisms underlying inflammatory pain remains to be fully elucidated.

The transient receptor potential (TRP) channel superfamily is one of the most studied type of receptors/channels in the context of pain. These proteins function as molecular sensors of multiple stimuli, including changes in pH (Jordt et al., 2000), chemical agents (Bessac and Jordt, 2008), temperature (Tan and McNaughton, 2016; Vandewauw et al., 2018), osmolarity (Strotmann et al., 2000), and even essential biological trace elements (Hu et al., 2009). TRP channels are generally non-selective Ca^2+^-permeable channels (Julius, 2013), classified into ankyrin (TRPA), canonical (TRPC), melastatin (TRPM), mucolipin (TRPML), polycystin (TRPP), and vanilloid (TRPV) subtypes (Wu et al., 2010). Notably, TRP channels play an important role in inflammatory pain. For example, TRPV1 is required for the development of inflammatory heat hyperalgesia (Caterina et al., 2000; Davis et al., 2000) and several pro-inflammatory mediators have been shown to be potentiate the function of this channel leading to nociceptor hypersensitivity (Chuang et al., 2001). Similarly, TRPA1 plays an important role in inflammation-related hyperalgesia (Kwan et al., 2006; Lennertz et al., 2012; Petrus et al., 2007). Pro-inflammatory mediators can also enhance the function of TRPA1 (Wang et al., 2008) and some inflammatory molecules, such as bradykinin, cause direct activation (Bandell et al., 2004). Yet, the molecular mechanisms involved in the activation and potentiation of other receptors in the context of inflammatory pain requires further investigation.

In recent years, TRPM3 has emerged as an important mediator of inflammatory pain. This channel is expressed in a large subset of both mouse (∼70-80%) and human (∼50%) sensory neurons (Vangeel et al., 2020; Vriens et al., 2011). Activation of TRPM3 by its agonist pregnenolone sulphate evokes pain (Kelemen et al., 2021; Ueda et al., 2001; Vriens et al., 2011) and its pharmacological inhibition has analgesic effects (Krügel et al., 2017; Straub et al., 2013). Of note, the expression and function of TRPM3 has been shown to increase during skin or bladder inflammation (Mulier et al., 2020; Zhao et al., 2022). However, the pro-inflammatory mediators and underlying mechanisms driving increased TRPM3 function remain poorly characterised. To answer this question, we first explored the expression and function of TRPM3 in mouse sensory neurons, before assessing the role of various pro-inflammatory mediators in modulating TRPM3 activity and expression. Finally, we studied the role of TRPM3 in inflammatory pain in a model of acute knee inflammation.

## 2. Material and Methods

### 2.1. Animals

All mouse experiments were performed in accordance with the Animals (Scientific Procedures) Act 1986 Amendment Regulations 2012 under Project Licenses granted to E.St.J.S. (PP5814995) by the Home Office and approved by the University of Cambridge Animal Welfare Ethical Review Body. Experiments were performed using a mixture of male and female C57BL6/J mice (8-14 weeks old). Mice were purchased from Envigo and housed conventionally with nesting material and a red plastic shelter in a temperature-controlled room at ∼21 °C, with a 12-hour light/dark cycle and access to food and water *ad libitum*.

### 2.2. Immunostaining of whole DRG

Mice were transcardially perfused with phosphate buffered saline (PBS) followed by 4% (w/v) paraformaldehyde (Sigma Aldrich) (in PBS, pH 7.4) under terminal anaesthesia (sodium pentobarbital, 200 mg/kg). DRG (L2-L5) were isolated and post-fixed in 4% (w/v) paraformaldehyde for 30 min, followed by overnight incubation in 30% (w/v) sucrose at 4 °C. Next, individual DRG were embedded in Shandon M-1 Embedding Matrix (Thermo Fisher Scientific), snap-frozen and stored at −80°C until processing. Cryosections (12 μm) were collated across Superfrost Plus slides (Thermo Fisher Scientific). Then, sections were washed with PBS-tween 20 (0.1%) and incubated in blocking buffer (PBS supplemented with 0.2% (v/v) Triton X-100, 5% (v/v) donkey serum and 1% (v/v) bovine serum albumin) for 2 hours at room temperature. Next, samples were incubated with primary antibodies: guinea pig anti-TRPV1 (1:500, Alomone, ACC-030-GP) and rabbit anti-TRPM3 (extracellular) (1:1000, Alomone, ACC-050) overnight at 4 °C. Slides were washed three times using PBS-tween 20 (0.1%) and incubated with species-specific conjugated secondary antibodies: donkey anti-rabbit Alexa Fluor 488 (1:1000, Invitrogen, A21206) and donkey anti-guinea pig Alexa Fluor-594 (1:1000, Jackson ImmunoResearch, 706-585-148) for 2 hours at room temperature. Slides were again washed three times using PBS-tween 20 (0.1%), mounted and imaged with an Olympus BX51 microscope and QImaging camera. Exposure levels were kept constant for each slide and the same contrast enhancements were made to all slides. Negative controls without the primary antibody were performed and showed no staining with either secondary.

For the analyses, sections were blinded for experimental groups and ImageJ software was used (version 1.53k). After manually selecting all neurons in a DRG section, we measured their mean grey value and normalised it between the highest and lowest intensity neurons in that section. The threshold for scoring a neuron as positive for a stain was set to the normalised minimum grey value across all sections plus 1.5 times the standard deviation (SD).

### 2.3. Drugs

The following drugs were used for Ca^2+^-imaging experiments: pregnenolone sulphate (PregS) (Sigma-Aldrich), capsaicin (Sigma-Aldrich), bradykinin (Tocris), recombinant murine interleukin (IL)-1β (PeproTech), recombinant murine IL-6 (PeproTech), recombinant murine tumour necrosis factor (TNF) α (PeproTech), and isosakuranetin (PhytoLab). Stock concentrations of PregS (50 mM and 500 mM) and isosakuranetin (5 mM and 10 mM) were dissolved in dimethyl sulfoxide (DMSO), capsaicin (1 mM) was dissolved in 100% ethanol, and bradykinin (1 mM), IL-1β (100 μg/mL), IL-6 (100 μg/mL) and TNFα (50 μg/mL) were dissolved in double distilled water (ddH_2_O). Drug aliquots were stored at -20°C until use.

For *in vivo* experiments (see below), isosakuranetin was dissolved in Miglyol 812 (kindly provided by IOI Oleo GmbH, Germany) containing 0.1% DMSO (Thermo Fisher Scientific). Mice were administered isosakuranetin intraperitoneally (2 mg/kg) (Aloi et al., 2023); Miglyol 812 containing 0.1% DMSO was used as a vehicle control.

### 2.4. Mouse DRG sensory neuron culture

Mouse DRG were bilaterally excised under a dissection microscope and cultured as previously described (Higham et al., 2024). In brief, after excision, isolated ganglia were incubated with Lebovitz L-15 Glutamax media (Invitrogen) containing 1 mg/mL type 1A collagenase (Sigma-Aldrich) and 6 mg/mL bovine serum albumin (BSA, Sigma-Aldrich) for 15 minutes (37°C, 5% CO_2_). Next, DRG were washed and subsequently incubated with L-15 media containing 1 mg/mL trypsin (Sigma-Aldrich) and 6 mg/mL BSA for 30 minutes (37°C, 5% CO_2_). Then, DRG were gently triturated using a P1000 pipette. Dissociated cells were resuspended in L-15 + GlutaMAX growth media supplemented with 10% (v/v) foetal bovine serum, 24 mM NaHCO_3_ 38 mM glucose and 2% (v/v) penicillin/streptomycin and plated on laminin- and poly-D-lysine-coated coverslips (MatTek). DRG neurons were incubated overnight at 37°C in 5% CO_2_ and were used for imaging after no more than 24 hours. When cells were incubated with inflammatory mediators, they were diluted in L-15 + GlutaMAX growth media supplemented with 10% (v/v) foetal bovine serum, 24 mM NaHCO_3_ 38 mM glucose and 2% (v/v) penicillin/streptomycin and incubated for 20 hours before Ca^2+^-imaging. For sensitisation experiments, DRG neurons were incubated with bradykinin 100 nM, IL-1β 10 ng/mL and TNFα 50 ng/mL, either in combination or separately. Vehicle-treated DRG neurons were incubated with L-15 + GlutaMAX growth media supplemented with 10% (v/v) foetal bovine serum, 24 mM NaHCO_3_ 38 mM glucose, 2% (v/v) penicillin/streptomycin and 0.1% ddH_2_O (v/v).

### 2.5. Ca^2+^ imaging of cultured DRG neurons

Following overnight seeding of DRG neurons, cells were loaded with 2 μM Fura-2AM (Invitrogen) for 20-30 min at 37 °C. Next, culture dishes were mounted on the stage of an inverted microscope (Olympus IX71) and visualised with a 20x objective (Olympus UApo/340). Cells were continuously superfused with Krebs solution at room temperature, containing (in mM): 150 NaCl, 6 KCl, 1 MgCl_2_, 1.5 CaCl_2_, 10 glucose, 10 4-(2-hydroxyethyl)-1-piperazineethanesulfonic acid (HEPES), and titrated to pH 7.4 with NaOH, during the entire length of the experiment through a flexible perfusion pencil (Automate Scientific) placed adjacent to the field of view. Superfusion was controlled with a clasp-valve, gravity-fed perfusion system (Biologic RSC-200). Neurons were illuminated alternately with a 340 nm and 380 nm LED to excite Fura2 (Cairn Research), and emission at 520 nm was imaged with a Hamamatsu OrcaFlash 4.0LT camera (1 fps, 100 ms exposure) and recorded using MetaFluor (Molecular Devices). The baseline intracellular Ca^2+^ concentration was monitored for 120 s before adding any other stimuli and a Krebs-based solution in which the KCl concentration was increased to 45 mM by iso-osmotic substitution of NaCl was used at the end of each experiment to identify DRG neurons. Agonists and mediators (see below) were diluted in the aforementioned Krebs solution at the final concentrations required.

Initial analysis was carried out in MetaFluor. Regions of interest (ROI) were manually traced around individual neurons, and the average emission intensity (minus background) at 520 nm per ROI was calculated following alternate illumination at 340 nm and 380 nm. The ratio of these intensities (F_340_/F_380_) was used as an indicator of intracellular [Ca^2+^]. A positive neuronal response to a given stimulus was determined if an increase in F_340_/F_380_ of >10% over baseline was measured. The magnitude of the response (peak minus baseline ratio, ΔF_340_/F_380_) to each stimulus was also measured. Cells not responding to 45 mM KCl with an abrupt F_340_/F_380_ increase were considered unhealthy or non-neuronal cells and hence excluded from further analysis.

### 2.6. Immunostaining of cultured DRG neurons

DRG neurons were isolated as in 2.4, but with neurons plated on Corning BioCoat Poly-D-Lysine/Laminin glass coverslips (Merck). After overnight incubation in supplemented L-15 + GlutaMAX growth media (see section 2.4), neurons were fixed with 4% (w/v) paraformaldehyde for 10 min, followed by 3 washes with PBS-tween 20 (0.1%). Next, coverslips were incubated in blocking buffer (PBS supplemented with 0.2% (v/v) Triton X-100, 5% (v/v) donkey serum and 1% (v/v) bovine serum albumin) for 2 hours at room temperature, followed by overnight incubation at 4 °C with primary antibodies: guinea pig anti-TRPV1 (1:500, Alomone, ACC-030-GP) and rabbit anti-TRPM3 (1:200, Alomone, ACC-050). Coverslips were washed thereafter three times using PBS-tween 20 (0.1%) and incubated with species-specific conjugated secondary antibodies: donkey anti-rabbit Alexa Fluor 488 (Invitrogen) and donkey anti-guinea pig Alexa Fluor-594 (Jackson ImmunoResearch) for 2 hours at room temperature. Coverslips were again washed three times using PBS-tween 20 (0.1%). During the second wash, coverslips were incubated with Hoechst 33342 (5 μg/mL, Tocris) for nuclei staining.

Imaging and subsequent image analysis were performed by a researcher blinded to the treatment conditions. Fluorescence images were captured with a Leica Stellaris 5 confocal microscope equipped with a 63X oil immersion objective (NA 1.4). A 3X digital zoom was applied, and the pinhole set to 1 Airy Unit to optimise optical sectioning. TRPV1-positive neurons were identified visually based on their fluorescence intensity. The z-axis height of each neuron was determined, and a single optical section was captured at the midpoint of the cell body. Image acquisition parameters, including laser power, detector gain, and pixel dwell time, were kept constant across all samples to ensure comparability.

Marker expression and translocation image analysis was performed using ImageJ software, following previously described methods (Balemans et al., 2019) with minor modifications. The cell membrane was defined as the outermost 1 μm of the cell surface, while the intracellular region (cytoplasm) was defined as the area 1 μm inward from the cell surface, excluding the nucleus (Supplementary Figure 3a). To generate the membrane ROI, the fluorescence image was first binarized. A median filter with a 3-pixel radius and the ’fill holes’ function was then applied to smooth and connect the thresholded regions. Subsequently, 8 pixels (equivalent to 0.96 μm) were eroded from the smoothed binary image. The membrane ROI was obtained by subtracting this eroded binary image from the original smoothed binary image. The intracellular ROI was created by subtracting the membrane ROI from the smoothed binary image of the whole cell. Mean fluorescence intensity was measured for both membrane and intracellular ROIs and the ratio of the membrane fluorescence to intracellular fluorescence was calculated. Translocation was calculated as the ratio of the mean fluorescence intensity of the membrane and the cytoplasm.

### 2.7. Knee joint intra-articular injections

All knee injections were performed through the patellar tendon and conducted under complete anaesthesia (100 mg/kg ketamine and 10 mg/kg xylazine) as described previously (Chakrabarti et al., 2020, 2018). For retrograde labelling of knee-innervating sensory neurons, 1.5 μL Fast Blue (2% in 0.9% saline; Polysciences) was injected in both knees one week before isolation. In animals undergoing knee inflammation with complete Freund’s adjuvant (CFA, 100 μg in 10 μL; Chondrex), Fast Blue was injected one week before CFA injection. CFA was also intra-articularly injected in one knee (ipsilateral knee was determined randomly) and the contralateral knee was used as internal control. A digital calliper was used to measure knee width before and 24 hours after CFA injection. To measure knee inflammation, the ratio of the width of the ipsilateral knee to that of the contralateral knee was calculated. Behavioural tests (digging and dynamic weight bearing, see below) were performed before (baseline) and 24 hours after CFA injection.

### 2.8. Digging behaviour

Digging behaviour was conducted as previously described (Pattison et al., 2024), mice being individually transferred to standard cages (49 x 10 x 12 cm) containing ∼3 cm of tightly packed fine-grain aspen midi 8/20 wood chip bedding (LBS Biotechnology). Mice were allowed 3 minutes to explore cages without interference and the behaviour was video recorded (iPhone 11 camera, Apple). Training sessions were carried out the day before baseline behaviours were assessed to allow mice to gain familiarity with the setup. Test sessions, also lasting 3 minutes in total, were performed before the intra-articular injection with CFA and 24 hours after. In mice treated with isosakuranetin, this was administered 24 hours after CFA injection and 30 minutes before the behavioural test. The number of visible burrows, defined as crater-like sites with displaced bedding material, remaining at the end of the 3-minute test was recorded. Next, digging behaviour was analysed offline in a blinded fashion. The latency of mice to begin digging was measured and the total time mice spent digging was scored independently by 2 observers, and the average digging times are reported (R^2^ = 0.80 ± 0.12 from 66 animals scored). Digging behaviour was defined as previously (Chakrabarti et al., 2020, 2018; Pattison et al., 2024), that is, by the vigorous disturbance of the digging substrate with all four limbs. Digging usually begins with the hind legs planted in a wide stance, followed by rapid movement of the substrate under the body using the forelimbs, and concludes with the hind limbs kicking the substrate backward to produce a burrow.

### 2.9. Dynamic weight bearing

Weight-bearing was measured in freely moving mice using an open field area with a pressure-sensitive floor (dynamic weight-bearing device, Bioseb) (Chakrabarti et al., 2020). Each mouse was tested for 3 minutes, of the 3-minute recording at least 1 minute was manually vadlidated for correct annotation of fore paw and hind paw prints. Only frames where paw identification was manually validated or assigned by the analysis software with the two highest confidence levels were taken forward for analysis. We measured the time that mice spent rearing (that is, standing on their hind legs only) in the arena, the time spent on individual rear paws, and the ratio of weight placed through contralateral vs. ipsilateral rear paws. Dynamic weight bearing was evaluated before the intra-articular injection with CFA and 24 hours after. In mice treated with isosakuranetin, this was administered 24 hours after CFA injection and 30 minutes before the behavioural test.

### 2.10. Statistical analyses

Normal (Gaussian) distribution was determined for all datasets using the Shapiro–Wilk normality test. Non-parametric tests were used for datasets where ≥1 group did not pass the normality test (assuming α = 0.05). Otherwise, parametric tests were used. The type of statistical test and the sample sizes for each experiment are provided in figure legends. Data were plotted and statistical analyses performed with Prism (GraphPad Software version 10.2.3). *P* values <0.05 were considered statistically significant.

## 3. Results

### 3.1. Functional expression of TRPM3 in murine DRG sensory neurons

Before examining the effects of inflammatory mediators on TRPM3 function, we first characterised the expression and function of TRPM3 in mouse DRG neurons under basal conditions using immunohistochemistry and Ca^2+^-imaging. In agreement with previous studies (Held et al., 2015; King et al., 2024; Vriens et al., 2011), we observed that ∼73% of DRG neurons display TRPM3 immunoreactivity (Figure 1a, b). Similarly, the TRPM3 agonist pregnenolone sulphate (PregS, 50 μM), an endogenous neurosteroid known to rapidly and reversibly activate TRPM3 (Wagner et al., 2008), elicited a robust increase in intracellular Ca^2+^ concentration ([Ca^2+^]_i_) increase in ∼76% of cultured mouse DRG neurons (Figure 1d, e) (IHC vs. Ca^2+^ imaging for TRPM3^+^ neuron identification: p = 0.061, Fisher’s exact test). Although TRPM3 immunoreactivity was prominently found in small- and medium-diameter neurons (<30 μm), the size distribution was similar to DRG neurons not expressing TRPM3 (Figure 1c). This contrasts with TRPV1 immunoreactivity, which was observed in ∼44% of DRG neurons (Supplementary figure 1a), which had a significantly smaller diameter than neurons devoid of TRPV1 (Supplementary figure 1b). Interestingly, most of the TRPV1-expressing DRG neurons also displayed TRPM3 immunoreactivity (∼82%, Figure 1f). In agreement with the expression analysis, using Ca^2+^-imaging, ∼87% of capsaicin-responding neurons also responded to PregS (Figure 1g, h) (IHC vs. Ca^2+^ imaging for TRPM3^+^-TRPV1^+^ neuron identification: p = 0.098, Fisher’s exact test). Altogether, these results confirm the high expression of TRPM3 in murine DRG neurons, and that this TRPM3 is highly expressed in in TRPV1-positive neurons, i.e. putative nociceptors.

**Figure 1.**
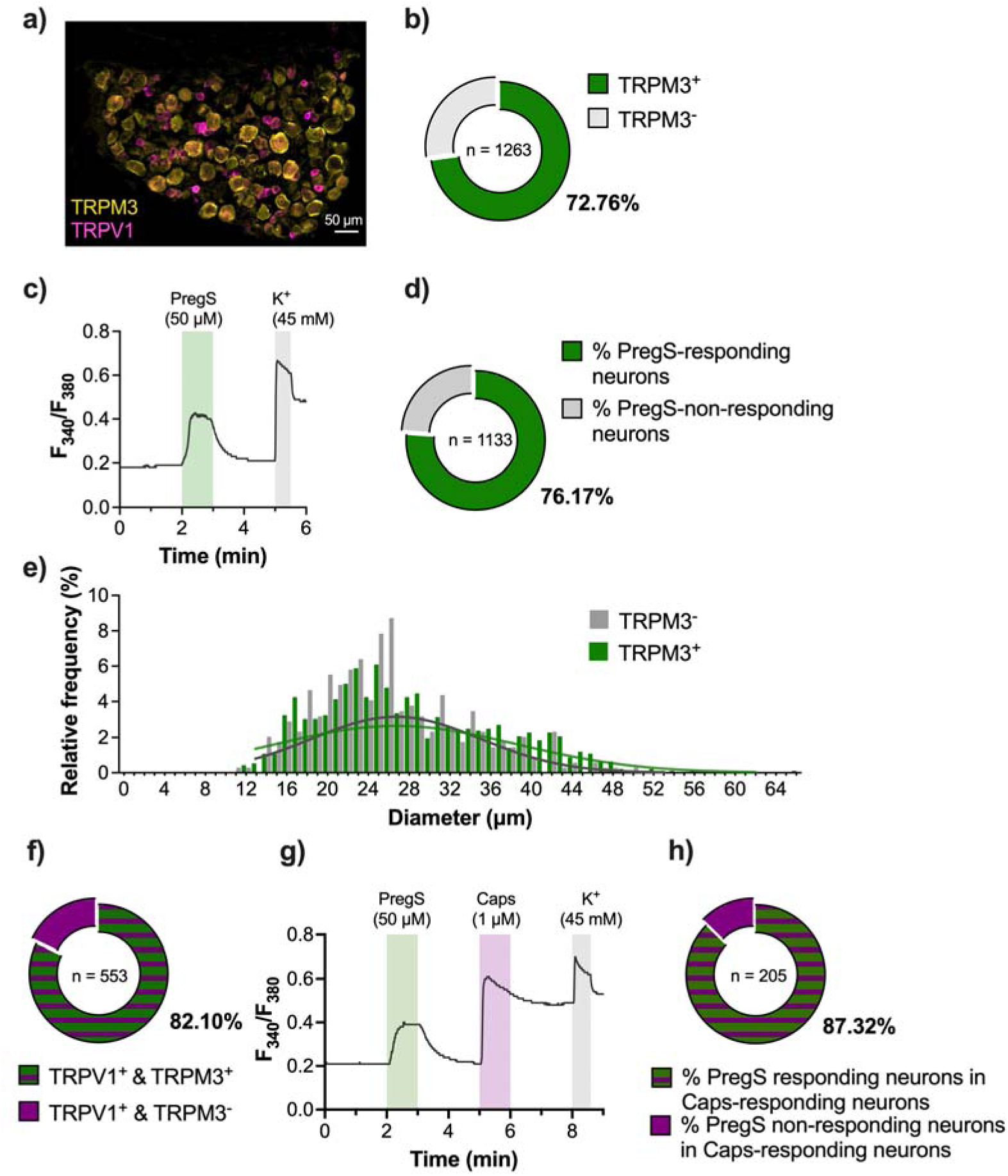
Functional expression of TRPM3 in murine DRG sensory neurons. **a)** Representative image of a whole DRG section (L4) showing TRPM3 (yellow) and TRPV1 (magenta) expression. **b)** Proportion of DRG neurons (L2-L5) that express TRPM3 in naïve mice (n = 1263 neurons, 8 sections, 2 mice). **c)** Representative Ca^2+^-imaging trace of cultured DRG neurons responding to the TRPM3 agonist pregnenolone sulphate (PregS). **d)** Proportion of cultured DRG neurons that respond to PregS (n = 1133 neurons, 3 mice). **e)** Relative frequency distribution of whole DRG neuron soma diameter expressing TRPM3 immunoreactivity. Least squares regression comparing best-fit values of mean soma size; R^2^ = 0.87 ± 0.008; degrees of freedom 9 (TRPM3^+^) and 7 (TRPM3^-^); F (1, 16) = 3.916; p = 0.0653. **f)** Proportion of DRG neurons (L2-L5) that express TRPV1 and TRPM3 in naïve mice (n = 553 neurons, 8 sections, 2 mice). **g)** Representative Ca^2+^ tracing of cultured DRG neurons responding to PregS and the TRPV1 agonist capsaicin (Caps). **h)** Proportion of cultured DRG neurons that respond to capsaicin and PregS (n = 205 neurons, 3 mice).

### 3.2. Pharmacological activation and inhibition of TRPM3 in murine DRG sensory neurons

Next, we sought to characterise the agonist concentration range required to activate TRPM3 in murine DRG neurons using Ca^2+^-imaging. Previous studies have shown that PregS elicits a concentration-dependent [Ca^2+^]_i_ increase in HEK293 cells heterologously expressing human and murine TRPM3 with EC_50_ values of 1 μM (Majeed et al., 2012, 2010) and 3 μM (Held et al., 2015), respectively. PregS also activates human stem cell-derived sensory neurons with an EC_50_ value of 48 μM (tested also using Ca^2+^-imaging) (Vangeel et al., 2020). In keeping with these findings, our assay yielded a dose-dependent [Ca^2+^]_i_ increase in murine DRG neurons with an EC_50_ of 0.677 μM (Figure 2a) and neuron activation frequency with an EC_50_ of 0.933 μM (Figure 2b). We observed similar activation frequencies and comparable Ca^2+^ responses between 50 μM and 500 μM of PregS (Figure 2b, c). These results suggest that 50 μM PregS is an adequate agonist concentration to confidently activate all TRPM3-expressing murine DRG neurons.

We next used the flavanone isosakuranetin to inhibit TRPM3 activity. This compound shows a marked specificity for TRPM3 compared to other TRP channels, such as TRPM8, TRPM7, TRPM1, TRPV1, and TRPA1 (Straub et al., 2013). We assessed whether isosakuranetin can not only prevent TRPM3-mediated Ca^2+^ responses when administered before PregS exposure, but also where it can abolish increases in DRG neuron [Ca^2+^]_i_ after TRPM3 activation. Application of isosakuranetin before PregS drastically reduced the number of PregS-responding neurons (Figure 2d, e) and the magnitude of the Ca^2+^ responses in TRPM3^+^ DRG neurons (Figure 2f). Similarly, isosakuranetin was able to block Ca^2+^ responses when applied during exposure to PregS (Figure 2g), eliciting an immediate and dramatic decrease in [Ca^2+^]_i_ (Figure 2h, i). These results are in agreement with published data (Held et al., 2015; Kelemen et al., 2021).

Altogether, our results indicate that PregS activates TRPM3 in murine DRG neurons with a similar potency to that observed following heterologous TRPM3 expression in HEK293 cells (Held et al., 2015). Furthermore, isosakuranetin abrogates TRPM3 activity when applied either before or after PregS stimulation.

**Figure 2.**
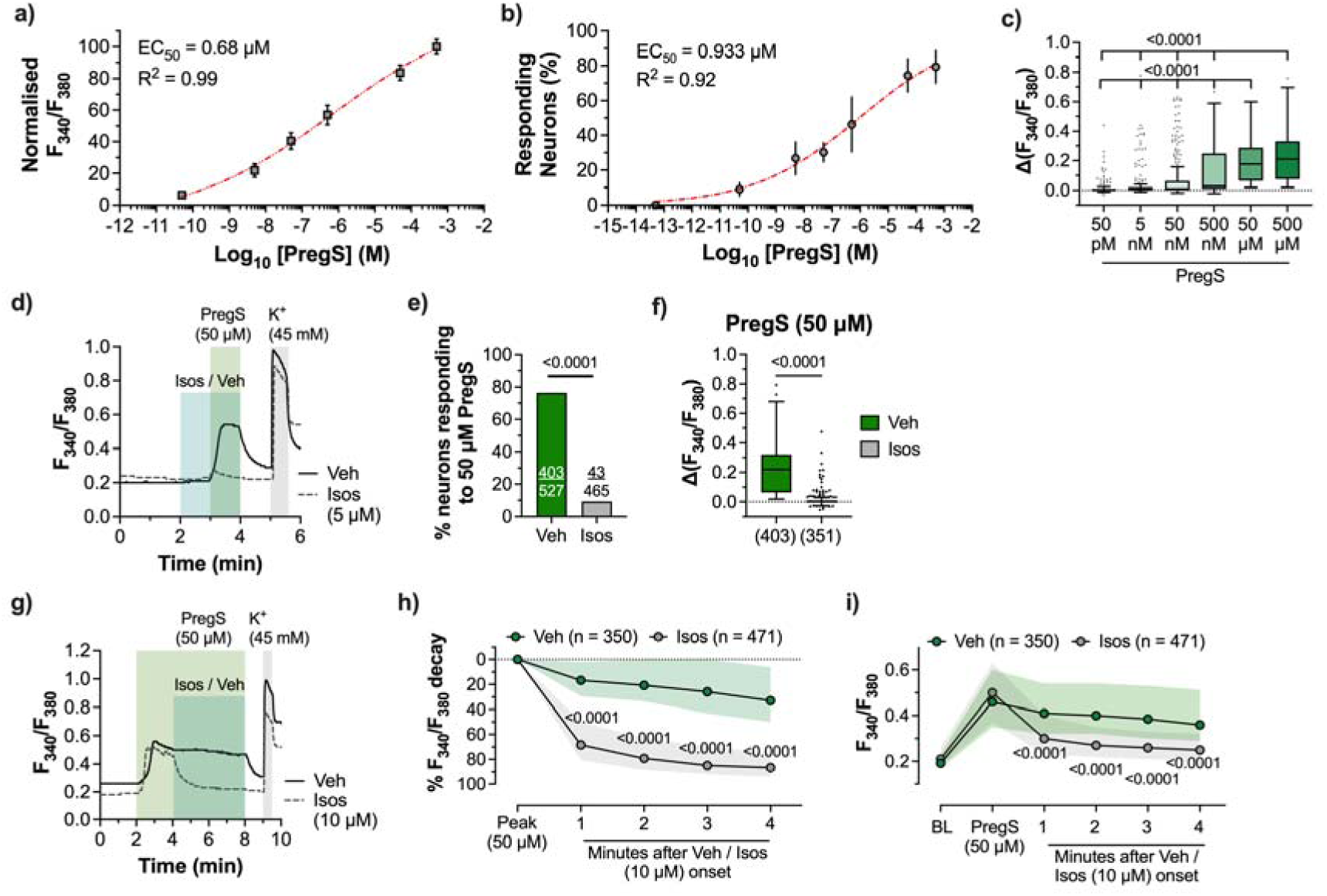
Pharmacological modulation of TRPM3 activation in murine DRG sensory neurons. **a)** Effect of increasing concentrations of PregS (n = 154-231 neurons) on Ca^2+^ responses in cultured DRG neurons. Absolute values were normalised using the maximum Ca^2+^ increase (ΔF_340_/F_380_) (data shown as mean ± SEM). **b)** Effect of increasing concentrations of PregS (n = 239-283 neurons) on the frequency of activation of DRG neurons (data shown as mean ± SD). **c)** Ratiometric [Ca^2+^]_i_ increase in cultured TRPM3^+^ DRG neurons (n = 154-231 neurons) in response to increasing concentrations of PregS (50 pM to 500 μM) (data shown as box and whiskers: centre line, median; box, 25^th^–75^th^ percentiles; whiskers, 10^th^–90^th^ percentiles; dots, outliers). Kruskal-Wallis test with Dunn’s multiple comparison test (p < 0.0001). **d)** Representative Ca^2+^-imaging trace of cultured DRG neurons responding to PregS after pre-exposure to isosakuranetin (Isos) or vehicle. **e)** Proportion of cultured DRG neurons that respond to PregS (veh, n = 527; isos, n = 465 neurons; 2 mice) following exposure to Isos or vehicle. Two-sided Fisher’s exact test. **f)** Ratiometric [Ca^2+^]_i_ increase in cultured TRPM3^+^ DRG neurons (veh, n = 403; isos, n = 351 neurons; 2 mice) in response to 50 μM PregS following exposure to Isos or vehicle (data shown as box and whiskers: centre line, median; box, 25^th^–75^th^ percentiles; whiskers, 10^th^–90^th^ percentiles; dots, outliers). Two-tailed Mann Whitney test. **g)** Representative Ca^2+^-imaging trace of cultured DRG neurons responding to PregS and subsequent application of Isos or vehicle. **h)** Proportion of ratiometric [Ca^2+^]_i_ decrease after application of Isos or vehicle in cultured DRG neurons that responded to PregS (veh, n = 350; isos, n = 471 neurons; 2 mice) (data shown as median ± IQR). Two-way repeated measures ANOVA with Sidak’s multiple comparisons test; F (4, 3252) = 103.0, P < 0.0001. **i)** Ratiometric [Ca^2+^]_i_ responses in DRG neurons responding to PregS and after application of Isos or vehicle (veh, n = 350; isos, n = 471 neurons; 2 mice) (data shown as median ± IQR). Two-way repeated measures ANOVA with Sidak’s multiple comparisons test; F (5, 4275) = 241.6, P < 0.0001. *P* values are shown in plots.

### 3.3. Functional effects of inflammatory mediators on TRPM3^+^ murine DRG sensory neurons

Several studies have shown that TRPM3 function is augmented in sensory neurons during inflammation (King et al., 2024; Mulier et al., 2020; Vanneste et al., 2022; Vriens et al., 2011; Zhao et al., 2022). However, little is known about the effects of specific inflammatory molecules in this process. To explore this, we first asked whether pro-inflammatory mediators activate neurons that functionally express TRPM3. To this end, we focused on several inflammatory molecules known to be elevated and play a role in complete Freund’s adjuvant (CFA)-induced inflammation, such as bradykinin (Davis and Perkins, 1996, 1994), IL-1β (Safieh-Garabedian et al., 1995; Woolf et al., 1997; Zucoloto et al., 2019), IL-6 (Ghasemlou et al., 2015; Zucoloto et al., 2019) and TNFα (Woolf et al., 1997; Zucoloto et al., 2019). All tested inflammatory mediators activated murine DRG neurons at the concentrations evaluated, although IL-6 elicited lower Ca^2+^ responses and in a significantly smaller fraction of neurons than other mediators tested (Supplementary Figure 2a-e). Among TRPM3^+^ DRG neurons, ∼36% of neurons also responded to bradykinin (Figure 3a, b), ∼34% responded to IL-1β (Figure 3d, e) and ∼38% responded to TNFα (Figure 3j, k); by contrast, only ∼4% of TRPM3^+^ DRG neurons responded to IL-6 (Figure 3g, h). The majority of DRG neurons responding to bradykinin, IL-1β and TNFα, also responded to 50 μM PregS: ∼86%, ∼82% and ∼92%, respectively (Figure 3c, f and l). This strongly suggests that most DRG neurons expressing receptors for these mediators co-express TRPM3. However, just over half of the DRG neurons responding to IL-6 (∼63%) also responded to PregS (Figure 3i), indicating that the prevalence of TRPM3 expression in these neurons is lower. We next evaluated whether pro-inflammatory mediator-responding DRG neurons display different sensitivity to PregS compared to pro-inflammatory mediator-insensitive neurons. While we observed no differences in the [Ca^2+^]_i_ increase to 50 μM PregS between bradykinin-, IL-1β-or IL-6-sensitive neurons and those insensitive to these mediators (Supplementary Figure 2f-h), TNFα-responding DRG neurons showed a slight, but statistically significant, increased sensitivity to 50 μM PregS (Supplementary Figure 2i).

Overall, these results demonstrate notable co-expression of TRPM3 and receptors for pro-inflammatory mediators in DRG neurons, including receptors of bradykinin, IL-1β and TNFα. This highlights the potential of these molecules to induce the TRPM3 sensitisation during inflammation. Moreover, our data suggest that TRPM3 might show an increased sensitivity in a subpopulation of DRG sensory neurons expressing receptors to TNFα.

**Figure 3.**
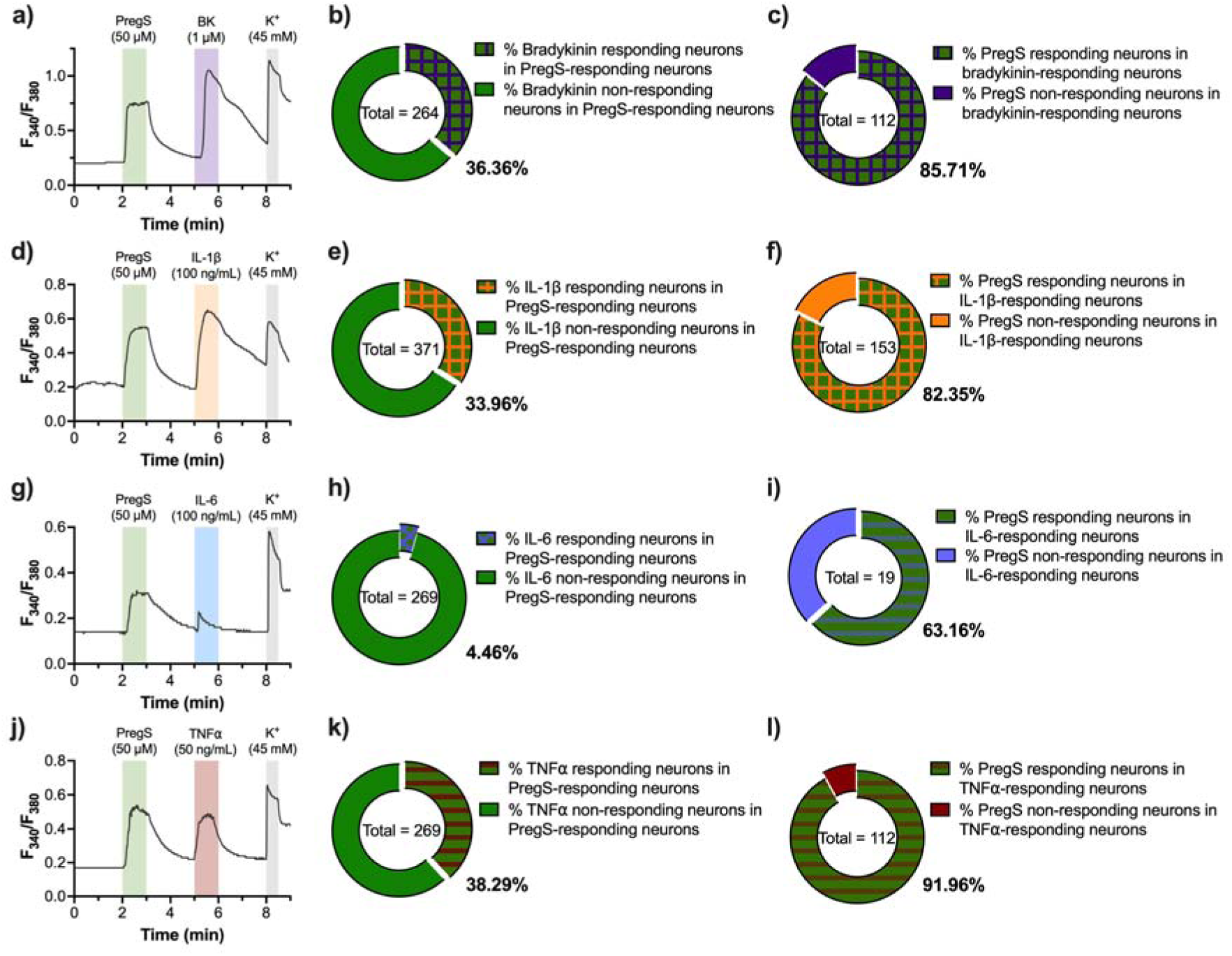
Responsiveness of TRPM3^+^ sensory neurons to pro-inflammatory mediators. **a)** Representative Ca^2+^-imaging trace of cultured TRPM3^+^ DRG neurons responding to bradykinin. **b)** Proportion of cultured TRPM3^+^ DRG neurons that respond to bradykinin (n = 264 neurons, 2 mice). **c)** Proportion of cultured bradykinin-responding DRG neurons that respond to PregS (n = 112 neurons, 2 mice). **d)** Representative Ca^2+^-imaging trace of cultured TRPM3^+^ DRG neurons responding to IL-1β. **e)** Proportion of cultured TRPM3^+^ DRG neurons that respond to IL-1β (n = 371 neurons, 2 mice). **f)** Proportion of cultured IL-1β-responding DRG neurons that respond to PregS (n = 153 neurons, 2 mice). **g)** Representative Ca^2+^-imaging trace of cultured TRPM3^+^ DRG neurons responding to IL-6. **h)** Proportion of cultured TRPM3^+^ DRG neurons that respond to IL-6 (n = 269 neurons, 2 mice). **f)** Proportion of cultured IL-6-responding DRG neurons that respond to PregS (n = 19 neurons, 2 mice). **j)** Representative Ca^2+^-imaging trace of cultured TRPM3^+^ DRG neurons responding to bradykinin. **h)** Proportion of cultured TRPM3^+^ DRG neurons that respond to bradykinin (n = 269 neurons, 2 mice). **f)** Proportion of cultured bradykinin-responding DRG neurons that respond to PregS (n = 112 neurons, 2 mice).

Given the low prevalence of TRPM3^+^ DRG neurons responding IL-6 (Figure 3g-i), we focused on the modulatory effects of the other mediators tested. Thus, we evaluated whether bradykinin, IL-1β, and TNFα can modulate TRPM3 function in murine DRG neurons. To this end, we used a low concentration of PregS, 50 nM, which we observed does not activate all TRPM3^+^ DRG neurons and elicits a weaker [Ca^2+^]_i_ increase compared to higher concentrations (Figure 2a-c). Moreover, we also used 50 μM PregS to identify TRPM3^+^ neurons, before characterising living neurons using KCl. To better represent the inflammatory conditions *in vivo* in our cultures, DRG neurons were incubated overnight with these mediators, rather than short-term incubation or acute administration. Overnight incubation with a cocktail of bradykinin (100 nM), IL-1β (10 ng/mL) and TNFα (50 ng/mL) enhanced DRG neuron Ca^2+^ responses to 50 nM PregS (Figure 4a) and the proportion of TRPM3^+^ neurons that responded to this low dose (Figure 4b). Furthermore, Ca^2+^ responses to stimulation with 50 μM PregS were also amplified by overnight incubation with the pro-inflammatory cocktail compared to vehicle incubation (Figure 4c), but the proportion of neurons responding to 50 μM PregS was not altered (Figure 4d).

We next investigated whether the potentiation effect of these mediators was due to their synergistic interaction or if any of them could individually enhance PregS-induced Ca^2+^ responses. Overnight incubation with bradykinin did not significantly increase the proportion of neurons responding to the low dose of PregS, nor did it amplify neuronal Ca^2+^ responses at either low or high concentrations of PregS (Figure 4e-g). By contrast, although IL-1β and TNFα did not increase the sensitivity of DRG neurons to a low dose of PregS or the proportion of responding cells, both cytokines did evoke increased [Ca^2+^]_i_ responses to a high dose of PregS compared to vehicle incubation (Figure 4i-k, m-o). Similar to the results with the three mediators combined, the proportion of DRG neurons activated by 50 μM PregS remained unchanged after individual overnight incubation with bradykinin, IL-1β, or TNFα (Figure 4h, l and p).

Altogether, these results demonstrate that inflammatory mediators can potentiate TRPM3 function in DRG neurons. IL-1β and TNFα, individually, were shown to potentiate Ca^2+^ responses to a high dose of PregS, unlike bradykinin, which had no effect. Notably, however, when all mediators were combined (as is more likely to occur during inflammation *in vivo*), they exerted a synergistic potentiation effect, increasing TRPM3 activity even in response to mild stimuli.

**Figure 4.**
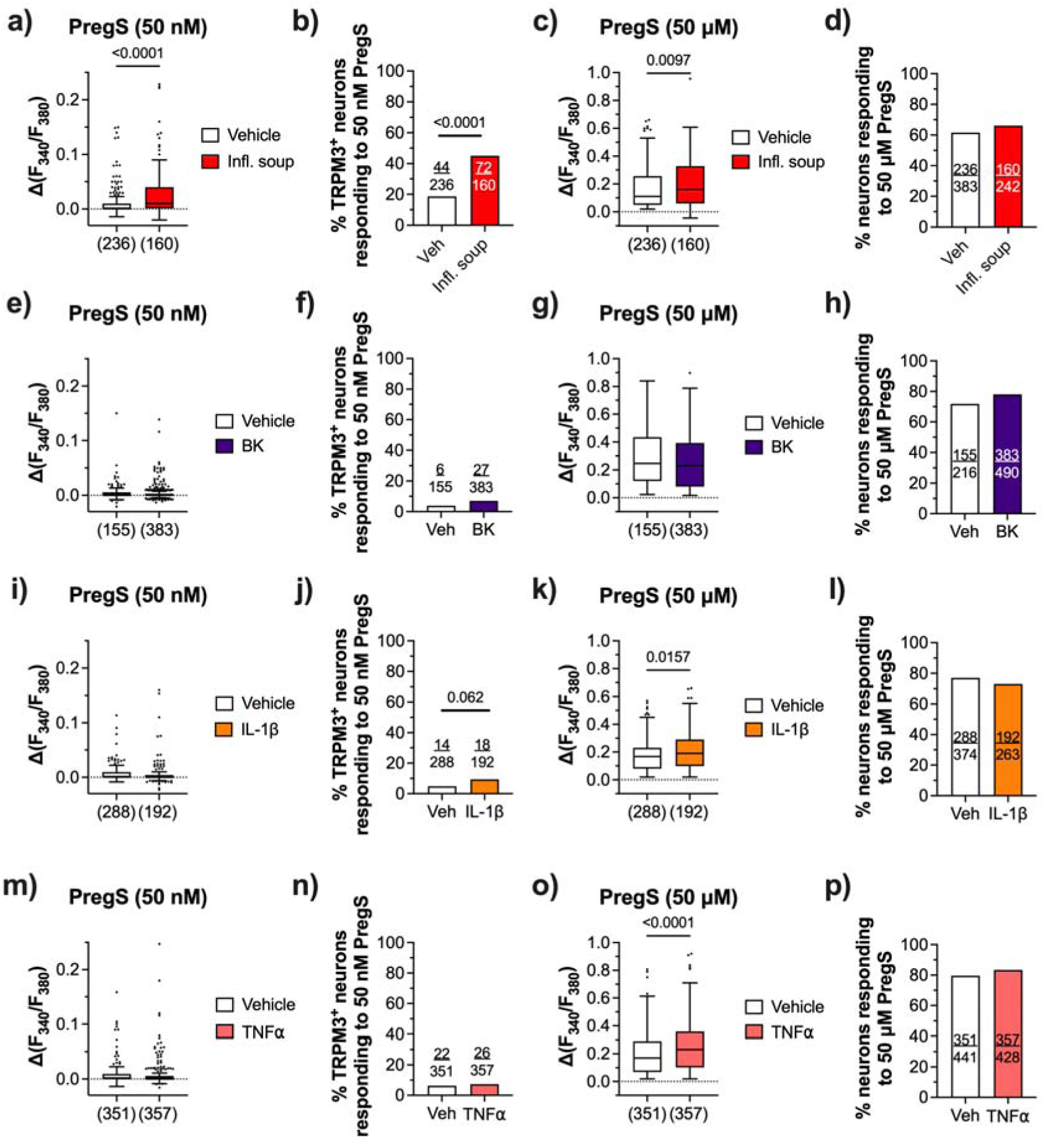
Potentiation/sensitisation of TRPM3^+^ by pro-inflammatory mediators. **a)** Ratiometric [Ca^2+^]i increase in cultured DRG neurons (vehicle n = 236 neurons; soup of inflammatory mediators, n = 160 neurons; 2 mice) in response to 50 nM PregS. Two-tailed Mann Whitney test. **b)** Proportion of cultured TRPM3^+^ DRG neurons that respond to 50 nM PregS (vehicle n = 236 neurons; soup of inflammatory mediators, n = 160 neurons; 2 mice). Two-sided Fisher’s exact test. **c)** Ratiometric [Ca^2+^]_i_ increase in cultured DRG neurons (vehicle n = 236 neurons; soup of inflammatory mediators, n = 160 neurons; 2 mice) in response to 50 μM PregS. Two-tailed Mann Whitney test. **d)** Proportion of cultured TRPM3^+^ DRG neurons that respond to 50 μM PregS (vehicle n = 383 neurons; soup of inflammatory mediators, n = 242 neurons; 2 mice). Two-sided Fisher’s exact test. **e)** Ratiometric [Ca^2+^]_i_ increase in cultured DRG neurons (vehicle n = 155 neurons; bradykinin (BK), n = 383 neurons; 2 mice) in response to 50 nM PregS. Two-tailed Mann Whitney test. **f)** Proportion of cultured TRPM3^+^ DRG neurons that respond to 50 nM PregS (vehicle n = 155 neurons; BK, n = 383 neurons; 2 mice). Two-sided Fisher’s exact test. **g)** Ratiometric [Ca^2+^]i increase in cultured DRG neurons (vehicle n = 155 neurons; BK, n = 383 neurons; 2 mice) in response to 50 μM PregS. Two-tailed Mann Whitney test. **h)** Proportion of cultured TRPM3^+^ DRG neurons that respond to 50 μM PregS (vehicle n = 216 neurons; BK, n = 490 neurons; 2 mice). Two-sided Fisher’s exact test. **i)** Ratiometric [Ca^2+^]_i_ increase in cultured DRG neurons (vehicle n = 288 neurons; IL-1β, n = 192 neurons; 2 mice) in response to 50 nM PregS. Two-tailed Mann Whitney test. **j)** Proportion of cultured TRPM3^+^ DRG neurons that respond to 50 nM PregS (vehicle n = 288 neurons; IL-1β, n = 192 neurons; 2 mice). Two-sided Fisher’s exact test. **k)** Ratiometric [Ca^2+^]_i_ increase in cultured DRG neurons (vehicle n = 288 neurons; IL-1β, n = 192 neurons; 2 mice) in response to 50 μM PregS. Two-tailed Mann Whitney test. **l)** Proportion of cultured TRPM3^+^ DRG neurons that respond to 50 μM PregS (vehicle n = 374 neurons; IL-1β, n = 263 neurons; 2 mice). Two-sided Fisher’s exact test. **m)** Ratiometric [Ca^2+^]_i_ increase in cultured DRG neurons (vehicle n = 351 neurons; TNFα, n = 357 neurons; 2 mice) in response to 50 nM PregS. Two-tailed Mann Whitney test. **n)** Proportion of cultured TRPM3^+^ DRG neurons that respond to 50 nM PregS (vehicle n = 351 neurons; TNFα, n = 357 neurons; 2 mice). Two-sided Fisher’s exact test. **o)** Ratiometric [Ca^2+^]_i_ increase in cultured DRG neurons (vehicle n = 351 neurons; TNFα, n = 357 neurons; 2 mice) in response to 50 μM PregS. Two-tailed Mann Whitney test. **p)** Proportion of cultured TRPM3^+^ DRG neurons that respond to 50 μM PregS (vehicle n = 441 neurons; TNFα, n = 428 neurons; 2 mice). Two-sided Fisher’s exact test. P values are shown in plots. In a, c, e, g, i, k, m and o, data shown as box and whiskers (centre line, median; box, 25^th^–75^th^ percentiles; whiskers, 10^th^–90^th^ percentiles; dots, outliers). Infl. soup, inflammatory soup.

### 3.4. Inflammatory mediators potentiate TRPM3 function in mouse DRG neurons without altering its protein expression or its translocation to the plasma membrane

Next, we investigated whether the increased TRPM3 functionality upon exposure to pro-inflammatory mediators results from upregulated expression, increased translocation to the plasma membrane, or increased sensitivity to stimuli (channel sensitisation). To this end, we incubated cultured murine DRG neurons with bradykinin, IL-1β, and TNFα, individually or in combination, and quantified the expression and cellular localisation of TRPM3 (Figure 5a, c and Supplementary Figure 3a). The expression and localisation of TRPV1 was also evaluated (Figure 5b, c). Overnight exposure to inflammatory mediators did not alter TRPM3 expression, either at the plasma membrane or in the cytoplasm (Figure 5d, e), nor did it affect the translocation of this channel to the membrane compared to exposure to the vehicle (Figure 5f). Although the lack of difference in TRPM3 expression remained when we compared TRPV1-expressing and TRPV1-lacking neurons (i.e. putative nociceptors vs. non-nociceptors, respectively), the overall expression of TRPM3 was higher in the TRPV1^+^ subpopulation (Supplementary Figure 3b-d). On the other hand, inflammatory mediators increased TRPV1 protein expression in both the plasma membrane and cytoplasm, but only when all three cytokines were present together, and not when incubated individually (Figure 5g, h).

These results strongly suggest that the enhanced function of TRPM3 upon exposure to the cocktail of bradykinin, IL-1β and TNFα is due to changes in the channel’s gating sensitivity to noxious stimuli, rather than upregulated protein expression or increased translocation to the plasma membrane of sensory neurons. In contrast, the overall protein expression of TRPV1 is increased in sensory neurons upon exposure to a pro-inflammatory environment, but not when exposed individually to bradykinin, IL-1β or TNFα.

**Figure 5.**
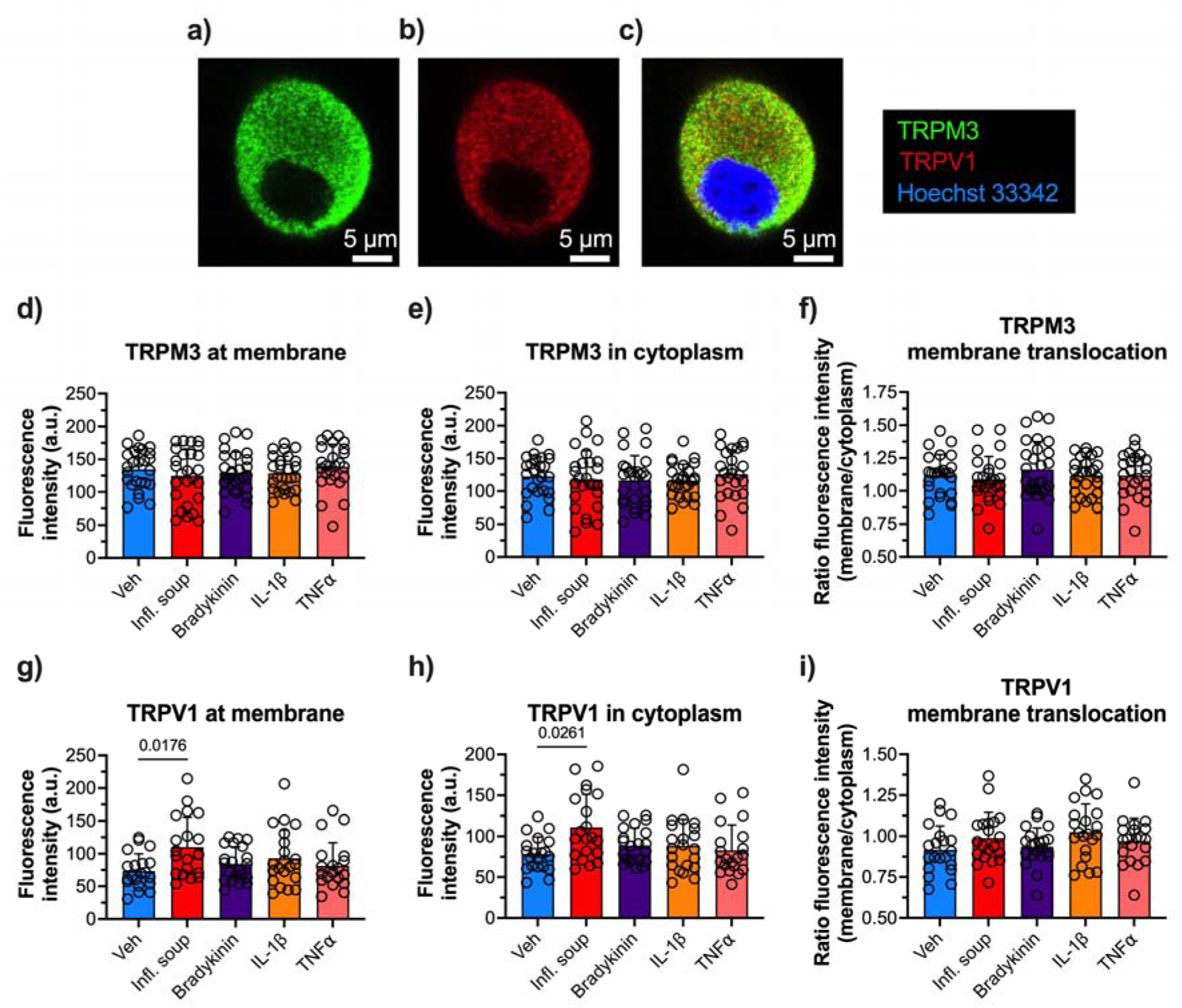
Inflammatory mediators increase expression and translocation of TRPV1, but not TRPM3, in DRG neurons. **(a-c)** Representative images of neuronal staining to visualise TRPM3, TRPV1 and the nucleus. Quantification of TRPM3 plasma membrane **(d**, Kruskal-Wallis test, P = 0.6203) and cytoplasm **(e**, one-way ANOVA F (4, 124) = 0.3676, P = 0.8313) expression in DRG neurons incubated with inflammatory mediators, measured as mean grey value. **f)** Quantification of TRPM3 translocation to the plasma membrane, measured as ratio between plasma membrane and cytoplasm fluorescence. One-way ANOVA F (4, 124) = 0.6263, P = 0.6446. In **d-f**: vehicle, n = 25 neurons; soup of inflammatory mediators, n = 26 neurons; bradykinin, n = 27 neurons; IL-1β, n = 26 neurons; and TNFα, n = 25 neurons. Quantification of TRPV1 plasma membrane **(d**, Kruskal-Wallis test, P = 0.0530) and cytoplasm **(e**, Kruskal-Wallis test, P = 0.0471) expression in DRG neurons incubated with inflammatory mediators, measured as mean grey value. **f)** Quantification of TRPV1 translocation to the plasma membrane, measured as ratio between plasma membrane and cytoplasm fluorescence. One-way ANOVA F (4, 97) = 1.755, P = 0.1443. In **g-i**: vehicle, n = 20 neurons; soup of inflammatory mediators, n = 20 neurons; bradykinin, n = 21 neurons; IL-1β, n = 21 neurons; and TNFα, n = 20 neurons. *P* values are shown in plots. In d, e, f, g, h and i, data shown as mean ± SD. Infl. soup, inflammatory soup.

### 3.5. Lack of change in the frequency of DRG neurons expressing the TRPM3 protein in a murine model of acute knee inflammation

After demonstrating that specific inflammatory mediators sensitise TRPM3 in mouse DRG neurons, we next aimed to investigate the role of this cation channel in the context of inflammatory pain. To this end, we used the well-characterised mouse model of CFA (Gould, 2000) to induce knee inflammation (Chakrabarti et al., 2018) and focused on knee-innervating DRG sensory neurons.

Firstly, we evaluated the expression of TRPM3 in retrogradely-labelled knee neurons in naïve mice. In keeping with previous studies in our lab (Bohic et al., 2023; Chakrabarti et al., 2020), knee-projecting neurons were restricted to only ∼5% of lumbar DRG (L2 – L5) (Supplementary figure 4a, b). Of these, the fraction of knee-innervating neurons displaying TRPM3 immunoreactivity was similar to that of the general DRG neuronal population (knee-innervating neurons 66.15% vs. all DRG neurons 72.76%, p = 0.2558; Supplementary Figure 4a, c and Figure 1b). Similarly, TRPV1 immunoreactivity was observed in a similar proportion of sensory neurons projecting to the knee compared to the general population of DRG neurons (knee-innervating neurons 49.23% vs. all DRG neurons 43.78%, p = 0.4425; Supplementary Figure 4a, d, and Supplementary Figure 1b).

Next, mice were injected with CFA unilaterally in the knee and changes in TRPM3 expression were evaluated using immunohistochemistry. Twenty-four hours after CFA injection, inflammation was confirmed in the ipsilateral knee compared to the injection of saline (Supplementary Figure 5a). Digging behaviour was evaluated as a measure of spontaneous pain (Pattison et al., 2024). As expected, CFA-injected mice spent less time digging, dug fewer burrows, and showed increased latency to dig compared to mice that received saline injections in the knee (Supplementary Figure 5b-d), indicating CFA-induced spontaneous knee pain. Thereafter, DRG from these mice were isolated and processed for immunohistochemistry. The proportion of knee-innervating sensory neurons displaying TRPM3 immunoreactivity was similar between mice injected with CFA and saline (Figure 6a). In keeping with previous findings in our lab (Chakrabarti et al., 2018), the fraction of TRPV1 immunoreactive DRG neurons innervating the ipsilateral knee in CFA-injected mice was increased compared to those injected with saline (Supplementary Figure 5e). The proportion of retrogradely-labelled knee neurons (Fast Blue^+^) was comparable between CFA- and saline-injected mice (Supplementary Figure 5f).

Altogether, our results indicate that the expression of TRPM3 in knee-innervating neurons is similar to that in the general population of DRG neurons. Moreover, acute inflammation induced by intra-articular injection of CFA does not increase the frequency of TRPM3 expression in knee sensory neurons at the protein level. This contrasts with a previous report showing that CFA-induced inflammation increases both the proportion of neurons projecting to inflamed skin tissue that express *Trpm3* mRNA transcripts and the levels its gene expression (Mulier et al., 2020). An upregulation at both the mRNA and protein levels was observed in a model of cyclophosphamide-induced cystitis (Zhao et al., 2022), albeit expression was analysed one week after induction of inflammation, compared to 24-hour CFA model used here and by Mulier and colleagues (Mulier et al., 2020). This suggests that an increase in TRPM3 protein expression may require a prolonged duration of inflammation.

### 3.6. Pharmacological inhibition of TRPM3 shows no benefit in pain behaviour in a murine model of acute knee inflammation

Finally, we aimed to evaluate the nociceptive role of TRPM3 in CFA-induced knee inflammation. Although we did not observe differences in the number of TRPM3-expressing DRG neurons innervating the knee between the CFA and control groups (Figure 6a), we speculated that the inflammatory environment in the knee might sensitise TRPM3 in sensory neurons, as we observed *in vitro* (Figure 4a-d), and hence contribute to the development of inflammatory knee pain. To test this hypothesis, we examined whether blocking TRPM3 with the specific inhibitor isosakuranetin can alleviate CFA-induced knee joint pain. As expected, given that CFA-induced inflammation is non-neurogenic (Hylden et al., 1992), treatment with isosakuranetin did not reduce CFA-induced inflammation compared to vehicle-treated animals (Figure 6b). Furthermore, isosakuranetin-treated control animals (receiving knee injections with saline) did not show knee swelling (Figure 6b). Although mice treated with isosakuranetin showed an improvement in the latency to start digging 24-hours after CFA injection compared to baseline (Figure 6c), these animals still exhibited deficits in the amount of time spent digging and in the number of burrows dug, compared to baseline (Figure 6d, e). Digging behaviour in these mice was similar to that of CFA-injected mice treated with vehicle. The treatment with isosakuranetin had no detrimental effects on the digging behaviour in mice receiving saline knee injections (Figure 6c-e).

Despite the ineffectiveness of isosakuranetin in reversing the CFA-induced deficits in digging behaviour, we also asked whether this TRPM3 inhibitor could alleviate inflammatory pain using a different approach. To this end, we assessed the body weight distribution across the limbs of mice treated with isosakuranetin. As we have observed previously (Chakrabarti et al., 2020), CFA-injected mice placed more weight on the non-injected rear limb compared to the inflamed one (Figure 6f), indicating mechanical allodynia due to knee inflammation. Furthermore, CFA-induced inflammation reduced the time the mice bore weight on the ipsilateral paw (Figure 6g), without affecting the time they bore weight on the contralateral paw (Supplementary Figure 6). Moreover, similar to what we have observed in other inflammatory pain models (Bohic et al., 2023), CFA-induced inflammation decreased the time mice spent rearing, that is, standing on their hind paws (Figure 6h). None of these parameters were improved after the treatment with isosakuranetin in mice subjected to CFA knee injections (Figure 5f-h), nor did treatment with this TRPM3 inhibitor in mice receiving saline knee injections have any impact on weight bearing (Figure 6f-h).

Altogether, these results indicate that blockade of TRPM3 activity is insufficient to alleviate CFA-induced inflammatory spontaneous pain and mechanical allodynia, suggesting that other channels and receptors, such as TRPV1 (Chakrabarti et al., 2018), play a more important role.

**Figure 6.**
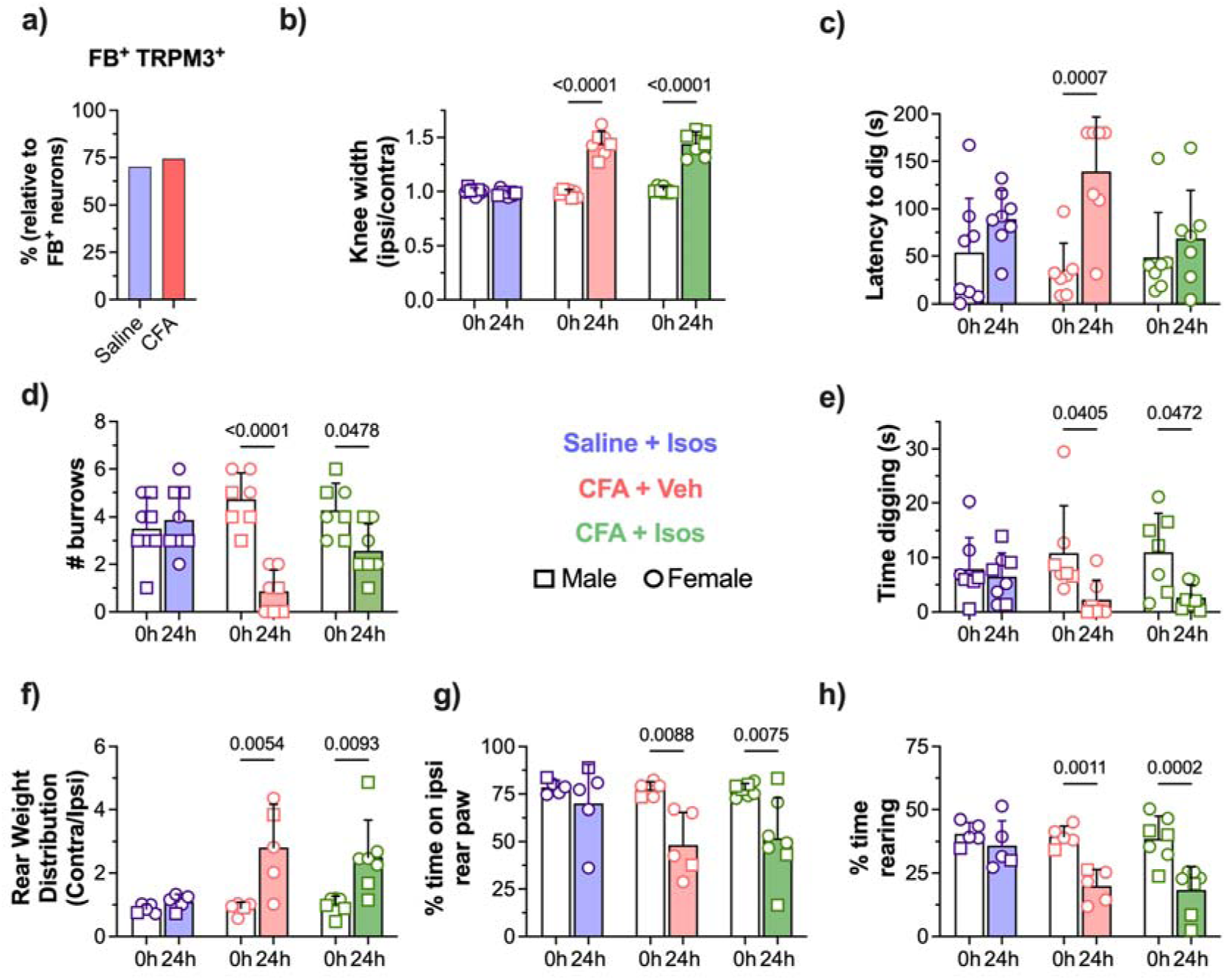
TRPM3 does not play a major role in the development of knee pain. **a)** Proportion of knee-innervating DRG neurons that express TRPM3 (saline n = 75/107 neurons [FB], 951 neurons [total]; CFA n = 58/78 neurons [FB], 998 neurons [total]; 3 mice/group). Two-sided Fisher’s exact test. **b)** Ratio of ipsilateral knee width to contralateral knee width on the day of injections and 24-hours after (saline+isos, n = 8; CFA+veh, n = 7; CFA+isos, n = 7). Two-way repeated measures ANOVA with Sidak’s multiple comparisons test; F (2, 19) = 53, P < 0.0001. **c)** Latency of mice to dig on the day of injections and 24-hours after (saline+isos, n = 8; CFA+veh, n = 7; CFA+isos, n = 7). Two-way repeated measures ANOVA with Sidak’s multiple comparisons test; F (2, 19) = 3.8, P < 0.0400. **d)** Time spent digging before injections and 24-hours after (saline+isos, n = 8; CFA+veh, n = 7; CFA+isos, n = 7). Two-way repeated measures ANOVA with Sidak’s multiple comparisons test; F (2, 19) = 1.8, P = 0.1920. **e)** Number of visible burrows at the conclusion of the 3-minute digging test on the day of injections and 24-hours after (saline+isos, n = 8; CFA+veh, n = 7; CFA+isos, n = 7). Two-way repeated measures ANOVA with Sidak’s multiple comparisons test; F (2, 19) = 11.32, P = 0.0006. **f)** Ratio of weight distribution between contralateral and ipsilateral paws during the 3-minute test on the day of injections and 24-hours after (saline+isos, n = 5; CFA+veh, n = 5; CFA+isos, n = 7). Two-way repeated measures ANOVA with Sidak’s multiple comparisons test; F (2, 14) = 3.1, P = 0.0006. **g)** Percentage of time spent standing on the ipsilateral rear paw at the conclusion of the 3-minute test on the day of injections and 24-hours after (saline+isos, n = 5; CFA+veh, n = 5; CFA+isos, n = 7). Two-way repeated measures ANOVA with Sidak’s multiple comparisons test; F (2, 14) = 1.8, P = 0.0003. **h)** Percentage of time spent rearing (standing on the hindpaws) during the 3-minute test on the day of injections and 24-hours after (saline+isos, n = 5; CFA+veh, n = 5; CFA+isos, n = 7). Two-way repeated measures ANOVA with Sidak’s multiple comparisons test; F (2, 14) = 4.7, P < 0.0001. *P* values are shown in plots. In b, c, d, e, f, g and h, data shown as mean ± SD.

## 4. Discussion

In this work, we assessed the impact of pro-inflammatory mediators on TRPM3 activity and its contribution to inflammatory pain. Inflammation generally results in an increased activity of ion channels and receptors in sensory neurons, which may lead to the development of aberrant nociceptive responses and pain. Notably, the role of TRP channels in the development of inflammatory pain is well recognised (Patapoutian et al., 2009). In recent years, the spotlight has turned to TRPM3, after several studies highlighted the role of this channel in inflammatory pain (King et al., 2024; Mulier et al., 2020; Vanneste et al., 2022; Vriens et al., 2011; Zhao et al., 2022). Here, we found that approximately one third of TRPM3-expressing DRG neurons in mice can be activated by pro-inflammatory mediators, including bradykinin, IL-1β an TNFα. In turn, most of the sensory neurons activated by these inflammatory mediators (∼85-90%) are activated by the TRPM3 agonist PregS. Of particular interest, we observed that overnight incubation with pro-inflammatory cytokines (IL1β and TNFα) can enhance TRPM3 function *in vitro*, an effect that was synergised when these molecules were combined with bradykinin. Additionally, we found that exposure to pro-inflammatory mediators does not alter the expression of this channel or its translocation to the cell membrane, suggesting that TRPM3 is sensitised through the activation of intracellular pathways that modulate its gating sensitivity to algogenic compounds. However, acute inhibition of TRPM3 *in vivo* was insufficient to alleviate spontaneous pain and mechanical allodynia in the CFA-induced knee inflammation model. Altogether, we demonstrated that pro-inflammatory mediators increase the functionality of TRPM3 in mouse DRG neurons through the sensitisation of the channel, which could consequently contribute to the development of pain states associated with the activation of the immune system.

We first characterised the expression and functionality of TRPM3 in murine DRG neurons. Utilising PregS as a TRPM3 activator, we found that 50 μM is sufficient to activate all TRPM3-expressing neurons *in vitro*. Moreover, the TRPM3 specific inhibitor isosakuranetin blocks the activity of this channel when present before or after the opening of TRPM3 by PregS, which suggests that it can be used for prophylactic or therapeutic approaches. Of note, isosakuranetin did not completely abolish PregS-induced DRG neuron activation. Although it has been shown that PregS does not act on other TRP channels, such as TRPV1, TRPV2, TRPA1 or TRPM8 (Chen and Wu, 2004; Vriens et al., 2011; Wagner et al., 2008), the small subset of PregS responding, isosakuranetin-insensitive DRG neurons implies the existence of other sensor(s) to this neurosteroid. However, *Trpm3*^-/-^ mice are resistant to PregS-induced pain when injected into the paw (Vriens et al., 2011), suggesting that the small fraction of PregS responding / isosakuranetin-insensitive sensory neurons has a negligible contribution to the development of pain following TRPM3 activation.

During inflammation, nociceptors undergo a process referred to as sensitisation, where they display increased responsiveness to their normal or subthreshold input. For example, the role of bradykinin on nociceptor sensitisation is well-established (Cesare and Mcnaughton, 1996; Dray and Perkins, 1993; Wang et al., 2006), specifically through its action on TRPV1 (Chuang et al., 2001; Lee et al., 2005; Mathivanan et al., 2016; Shin et al., 2002; Sugiura et al., 2002) and TRPA1 (Bautista et al., 2006; Brierley et al., 2009). In keeping with this, other pro-inflammatory molecules, such TNFα and IL-1β, have been shown to sensitise nociceptors and potentiate TRPV1 functionality (Ebbinghaus et al., 2012; Hu et al., 2010; Nicol et al., 1997; Prado et al., 2021). Although previous studies have indicated that TRPM3 function is potentiated in nociceptors innervating inflamed tissue (Mulier et al., 2020; Zhao et al., 2022), the inflammatory mediators causing this effect are poorly characterised. We found that TRPM3 function could be modestly enhanced by either IL-1β or TNFα when present individually, whereas no such effect was observed with bradykinin. The lack of bradykinin-induced TRPM3 sensitisation is somewhat surprising because recent findings have shown that responses to PregS were enhanced in neurons that had a preceding response to bradykinin (Behrendt et al., 2022). The authors of this study concluded that bradykinin may therefore contribute to the sensitisation of TRPM3 during inflammatory states (Behrendt et al., 2022). However, Behrendt and colleagues did not compare the effects of bradykinin with a vehicle or control. Instead, they examined the effects of a second application of PregS on neurons that either responded to bradykinin or failed to do so. Therefore, this study (Behrendt et al., 2022) primarily investigated the impact of bradykinin in TRPM3 tachyphylaxis, that is, channel desensitisation.

Notably, we observed that combining bradykinin, IL-1β, and TNFα, likely a better but still incomplete representation of what occurs during inflammatory states, enhanced Ca^2+^ responses in DRG neurons to both high and low doses of PregS. This indicates that the combination of various pro-inflammatory mediators has a synergistic potentiating effect. Nociceptor sensitisation may result from alterations in the gating properties of ion channels, including TRP channels, or from an increased abundance of these channels or other receptors in the plasma membrane (Linley et al., 2010). Interestingly, the mechanisms underlying nociceptor sensitisation may depend on the cellular context. Indeed, an elegant study showed that ATP sensitises nociceptors by two different mechanisms depending on the neuronal subtype (Devesa et al., 2014). In peptidergic neurons, ATP induces newly synthesised TRPV1 channels to move to the cell membrane, potentiating their excitability. However, in non-peptidergic neurons, ATP modulates the gating activity of TRPV1, causing hypersensitivity (Devesa et al., 2014). In the present study, we found that none of the mediators tested, neither individually nor in combination, increased the expression of TRPM3 or its translocation to the cell membrane. Although we did not specifically compare peptidergic and non-peptidergic neurons, we failed to detect differences in TRPM3 protein upregulation and translocation between TRPV1-expressing and non-expressing neurons. Interestingly, however, TRPV1-expressing sensory neurons displayed increased expression of TRPM3 compared to TRPV1-lacking neurons, suggesting that nociceptors might be more easily activated by TRPM3 agonists. Altogether, our results suggest that the increased responsiveness of TRPM3 under inflammatory conditions occurs through the modulation of the gating properties of this channel, rather than major increases in expression or trafficking to the cell membrane. Defining precise molecular pathways underlying this process will be the subject of future work. Moreover, the effect of other inflammatory mediators on TRPM3 expression and sensitisation remains to be elucidated.

Finally, we used the CFA-induced inflammatory pain model (Chakrabarti et al., 2018; Gould, 2000) to gain insights into the role of TRPM3 in the development of inflammatory pain *in vivo*. Injection of CFA causes mechanical allodynia in the inflamed area (Zhang et al., 2021), along with both mechanical and thermal hyperalgesia (Caterina et al., 2000; Hsieh et al., 2017; Safieh-Garabedian et al., 1995). This is associated with an increased fraction of neurons that express TRPV1 (Breese et al., 2005; Chakrabarti et al., 2018) and an increased expression of the TRPV1 channel itself (Yu et al., 2008; Zhang et al., 2005). Furthermore, we previously showed that injection of CFA into the knee joint impairs the digging behaviour in mice, an effect that can be reversed by pharmacological inhibition of TRPV1 (Chakrabarti et al., 2018), mouse digging being a relevant and measurable behaviour indicative of mouse well-being (Chakrabarti et al., 2020; Pattison et al., 2024). In the context of knee inflammation, deficits in digging can likely be attributed to pain caused by the associated aberrant immune response. Of interest, earlier work has revealed that inhibition of either only TRPV1 (Caterina et al., 2000; Davis et al., 2000; Gavva et al., 2005; Honore et al., 2005) or TRPM3 (Alkhatib et al., 2019; Straub et al., 2013; Vriens et al., 2011) is sufficient to fully abrogate inflammatory heat hyperalgesia in a variety of inflammatory pain models. However, in our hands, pharmacological inhibition of TRPM3 did not reverse CFA-induced digging deficits nor restore CFA-induced adjustments in body weight bearing. Therefore, our data suggest a less important role of TRPM3 in inflammatory spontaneous pain and inflammatory allodynia in this model. Although we cannot rule out a lack of target engagement by isosakuranetin, the dose used was similar to that used in other behavioural studies where effects were observed (Aloi et al., 2023; Su et al., 2021). However, it is important to note that the role of TRPM3 in pain hypersensitivity appears to depend on the nature of the inflammatory response. For example, a recent study found that TRPM3 plays a significant role in mechanical and thermal hyperalgesia, as well as bladder hyperreactivity, in a model of cyclophosphamide-induced bladder inflammation (Zhao et al., 2022). These results are in keeping with another study in which TRPM3 blockade alleviated mechanical hyperalgesia and cold hypersensitivity in a model of chemotherapy-induced peripheral neuropathic pain (Aloi et al., 2023). Conversely, however, another study demonstrated that inhibition of TRPM3 reduced heat hyperalgesia and spontaneous pain but did not affect mechanical or cold hyperalgesia in a neuropathic pain model following nerve injury (Su et al., 2021). Taken together, despite its well-established role in inflammatory heat hyperalgesia (Alkhatib et al., 2019; Straub et al., 2013; Vriens et al., 2011), the function of TRPM3 in mechanical hyperalgesia/allodynia and spontaneous pain seems to be determined by the associated immune signature. Characterising the specific and combined effects of various immune molecules on this channel is crucial for furthering our understanding of TRPM3’s relative contribution to pain signalling in different neuro-immune scenarios.

In conclusion, our study characterises the effect of various pro-inflammatory molecules, in isolation and in combination, on the modulation of TRPM3 function. Importantly, we demonstrate that the sensitising effect of these pro-inflammatory mediators is amplified when they are co-administered, resulting in TRPM3 being more strongly activated by lower agonist concentration. Furthermore, although pharmacological blockade of this channel did not significantly alleviate spontaneous or mechanical inflammatory pain in the model of CFA-induced knee inflammation, we propose that the nature of the immune response, along with the specific inflammatory mediators present in the microenvironment, likely determine the role of TRPM3 in modulating pain signalling. Further studies are needed to fully characterise the role of this channel in inflammatory and other types of pain.

## Acknowledgements

This work was principally funded by a Pain Relief Foundation research grant (J.A.-L. and E.St.J.S.). Furthermore, this work was partly supported by a joint and equal investment from UKRI and Versus Arthritis (MR/W002426/1) as part of the ADVANTAGE visceral pain consortium through the Advanced Pain Discovery Platform (APDP) initiative (J.A.L., L.A.P. and E.St.S.J.). T.K.L. acknowledges funding from UKRI Guarantee Marie Skłodowska-Curie Actions (MSCA) postdoctoral fellowship (EP/X023117/1). L.W.P. was funded by an AstraZeneca PhD studentship (G113502). Finally, the authors gratefully acknowledge the Cambridge Advanced Imaging Centre for their support and assistance in this work.

## Disclosures

No conflicts of interest, financial or otherwise, are declared by the authors.

## Author contributions

J.A.-L. conceived and designed research; J.A.-L., T.K.L., and L.A.P. performed experiments; J.A.-L., T.K.L., and L.W.P. analysed data; D.C.B. provided experimental support; J.A.-L. and E.St.J.S. interpreted results of experiments; J.A.-L. prepared figures; J.A.-L. drafted manuscript; J.A.-L. and E.St.J.S. edited and revised manuscript; J.A.-L., T.K.L., L.A.P., L.W.P., D.C.B., and E.St.J.S. approved final version of manuscript.

## Data availability

Data sets supporting the conclusions of this article are available in University of Cambridge Apollo Repository (<u>10.17863/CAM.111809</u>).

## Supplementary information

**Supplementary Figure 1.**
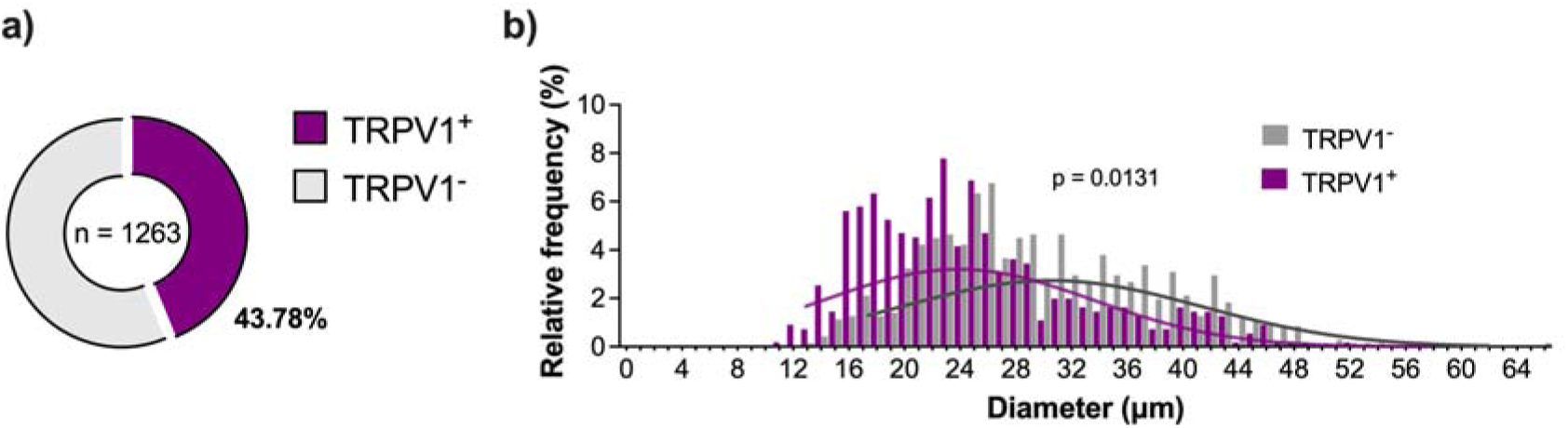
Expression of TRPV1 in murine DRG sensory neurons. **a)** Proportion of DRG neurons (L2-L5) that express TRPV1 in naïve mice (n = 1263 neurons, 8 sections, 2 mice). **b)** Relative frequency distribution of whole DRG neuron soma diameter expressing TRPV1 immunoreactivity. Least squares regression comparing best-fit values of mean soma size; R2 = 0.87 ± 0.008; degrees of freedom 8 (TRPV1^+^) and 8 (TRPV1^-^); F (1, 16) = 7.795; p = 0.0131. *P* values are shown in plots. Least squares regression comparing best-fit values of mean soma size. R^2^ = 0.77 ± 0.038.

**Supplementary Figure 2.**
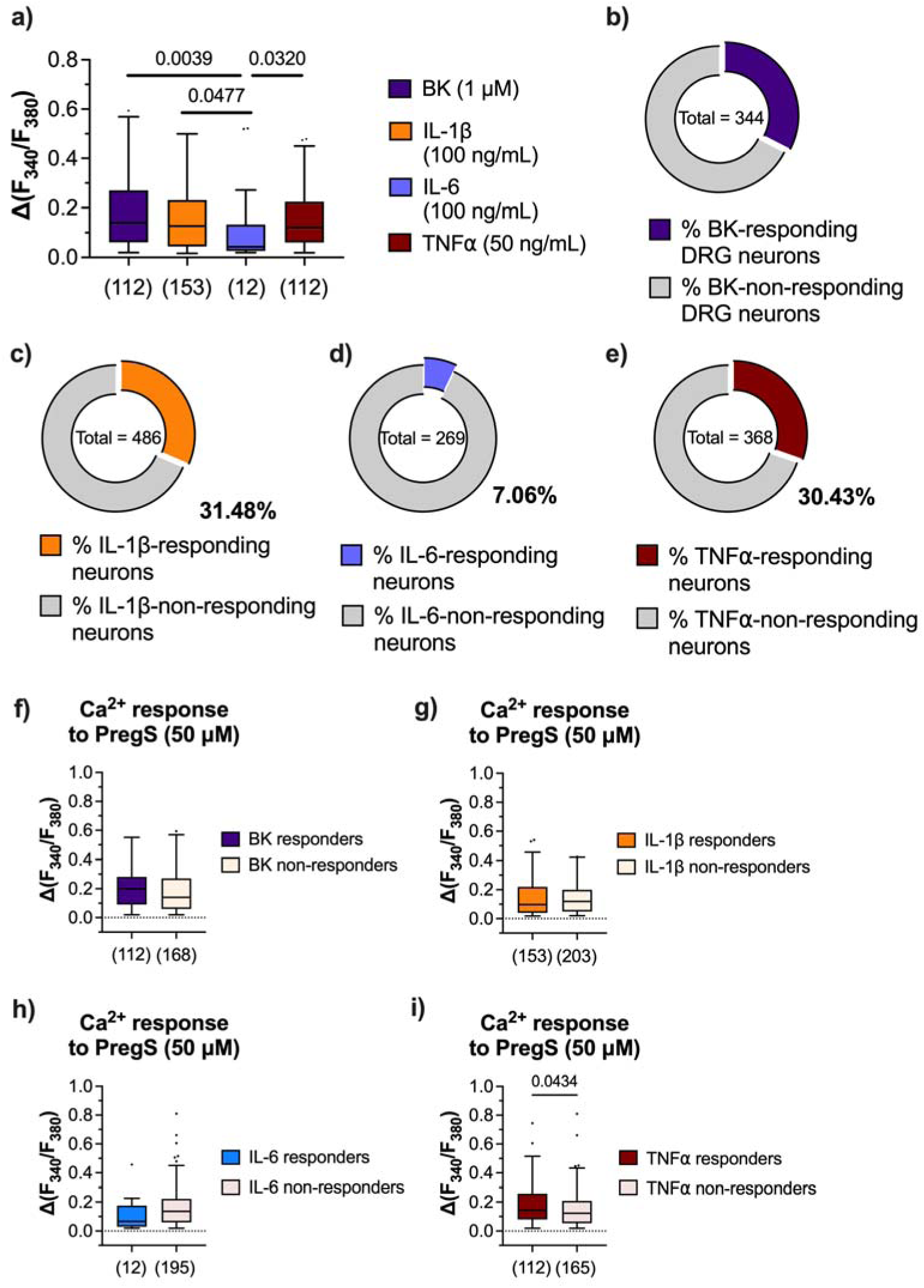
Responsiveness of TRPM3^+^ sensory neurons to pro-inflammatory mediators. **a)** Ratiometric [Ca^2+^]_i_ increase in cultured DRG neurons in response to bradykinin (n = 112; 2 mice), IL-1β (n = 153; 2 mice), IL-6 (n = 12; 2 mice) and TNFα (n = 112; 2 mice). Kruskal-Wallis test with Dunn’s multiple comparison test (p = 0.0082) **b)** Proportion of cultured DRG neurons that respond to bradykinin (n = 344 neurons, 2 mice). **c)** Proportion of cultured DRG neurons that respond to IL-1β (n = 486 neurons, 2 mice). **d)** Proportion of cultured DRG neurons that respond to IL-6 (n = 269 neurons, 2 mice). **e)** Proportion of cultured DRG neurons that respond to TNFα (n = 368 neurons, 2 mice). **f)** Ratiometric [Ca^2+^]_i_ increase in response to PregS in bradykinin-responding (n = 112, 2 mice) and non-responding (n = 168, 2 mice) cultured DRG neurons. Two-tailed Mann Whitney test. **g)** Ratiometric [Ca^2+^]_i_ increase in response to PregS in IL-1β-responding (n = 153, 2 mice) and non-responding (n = 203, 2 mice) cultured DRG neurons. Two-tailed Mann Whitney test. **h)** Ratiometric [Ca^2+^]i increase in response to PregS in IL-6-responding (n = 12, 2 mice) and non-responding (n = 195, 2 mice) cultured DRG neurons. Two-tailed Mann Whitney test. **i)** Ratiometric [Ca^2+^]_i_ increase in response to PregS in TNFα-responding (n = 112, 2 mice) and non-responding (n = 165, 2 mice) cultured DRG neurons. Two-tailed Mann Whitney test. *P* values are shown in plots. In a, f, g, h and i, data shown as box and whiskers (centre line, median; box, 25^th^–75^th^ percentiles; whiskers, 10^th^–90^th^ percentiles; dots, outliers).

**Supplementary Figure 3.**
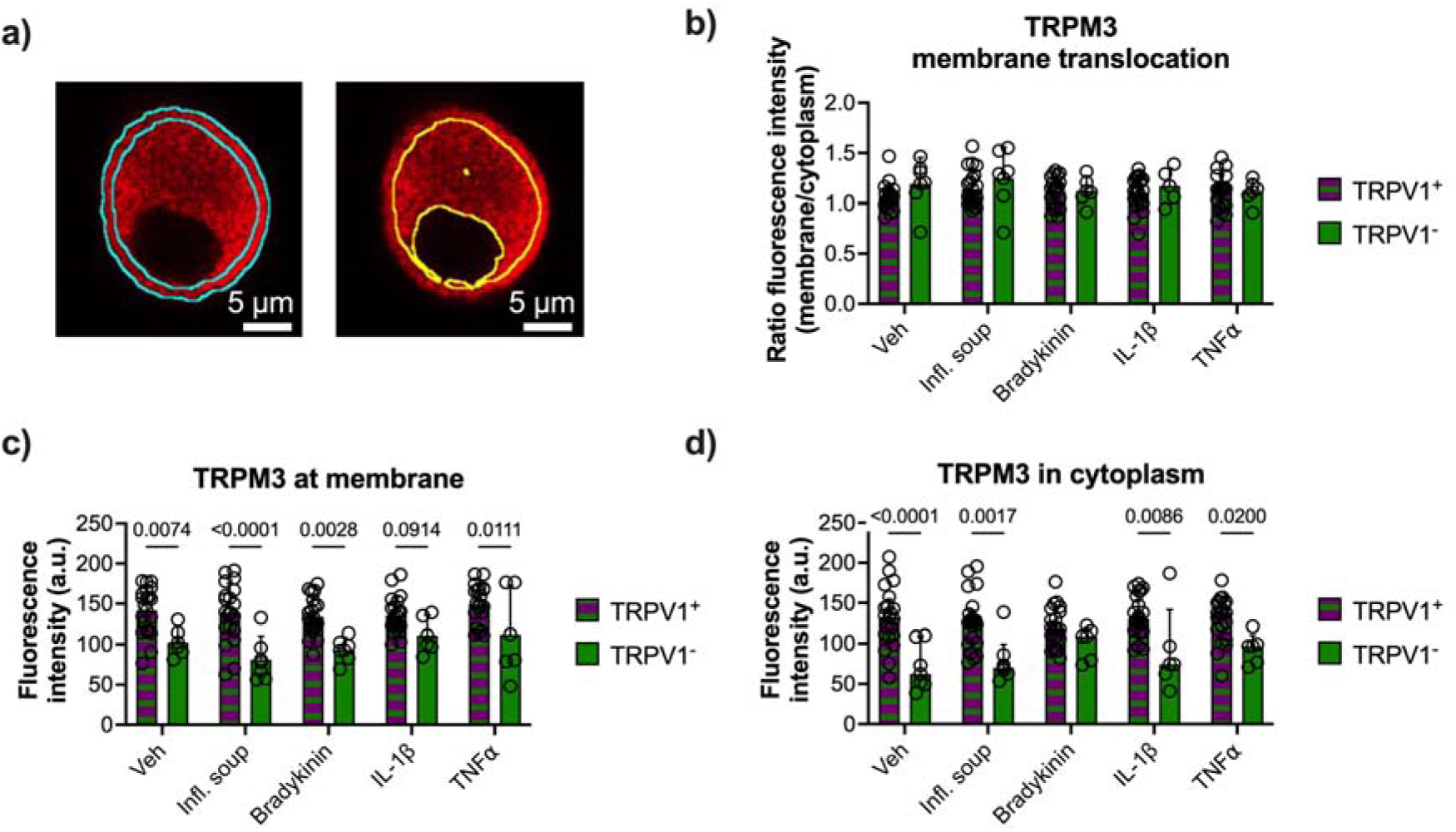
Inflammatory mediators do not affect the expression and translocation of TRPM3 in either TRPV1-expressing or non-TRPV1-expressing DRG neurons. **a)** Representative image of the cell area selected for channel expression quantification at the plasma membrane (blue lines, left), defined as outermost 1 μm of the cell surface, and the cytoplasm (yellow lines, right), defined as the area 1 μm inward from the cell surface, excluding the nucleus. **b)** Quantification of TRPM3 translocation to the plasma membrane, measured as ratio between fluorescence in the membrane and the cytoplasm in TRPV1^+^ and TRPV1^-^ DRG neurons. Two-way repeated measures ANOVA with Sidak’s multiple comparisons test; F (4, 119) = 0.5276, P = 0.7157. Quantification of TRPM3 expression in the plasma membrane **(c,** 2-way repeated measures ANOVA with Sidak’s multiple comparisons test; F (4, 119) = 40.59, P < 0.0001) and cytoplasm **(d,** 2-way repeated measures ANOVA with Sidak’s multiple comparisons test; F (4, 119) = 36.31, P < 0.0001) in TRPV1^+^ and TRPV1^-^ DRG neurons incubated with inflammatory cytokines, measured as mean grey value. In **b-d**: vehicle, n = 20 TRPV1^+^ neurons and n = 5 TRPV1^-^ neurons; soup of inflammatory mediators, n = 20 TRPV1^+^ neurons and n = 6 TRPV1^-^ neurons; bradykinin, n = 21 TRPV1^+^ neurons and n = 6 TRPV1^-^ neurons; IL-1β, n = 21 TRPV1^+^ neurons and n = 5 TRPV1^-^ neurons; and TNFα, n = 20 TRPV1^+^ neurons and n = 5 TRPV1^-^ neurons. *P* values are shown in plots. In b, c and d, data shown as mean ± SD.

**Supplementary Figure 4.**
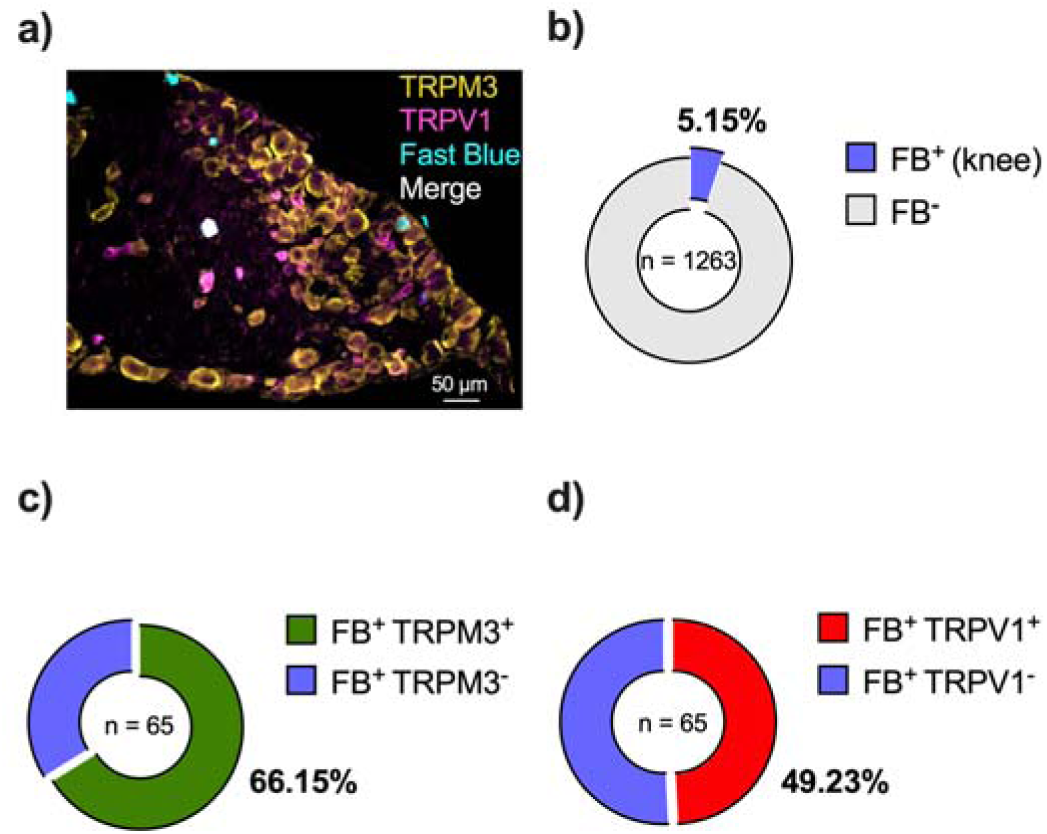
Expression of TRPM3 and TRPV1 in knee-innervating DRG neurons. **a)** Representative image of a whole DRG section (L4) showing TRPM3 (yellow), TRPV1 (magenta) and Fast Blue (cyan) expression. **b)** Proportion of Fast Blue^+^ sensory neurons after dye injection into the knee. **c)** Proportion of knee-innervating DRG neurons that express TRPM3 (n = 43/65 neurons, 1263 neurons [total]). **c)** Proportion of knee-innervating DRG neurons that express TRPV1 (n = 32/65 neurons, 1263 neurons [total]).

**Supplementary Figure 5.**
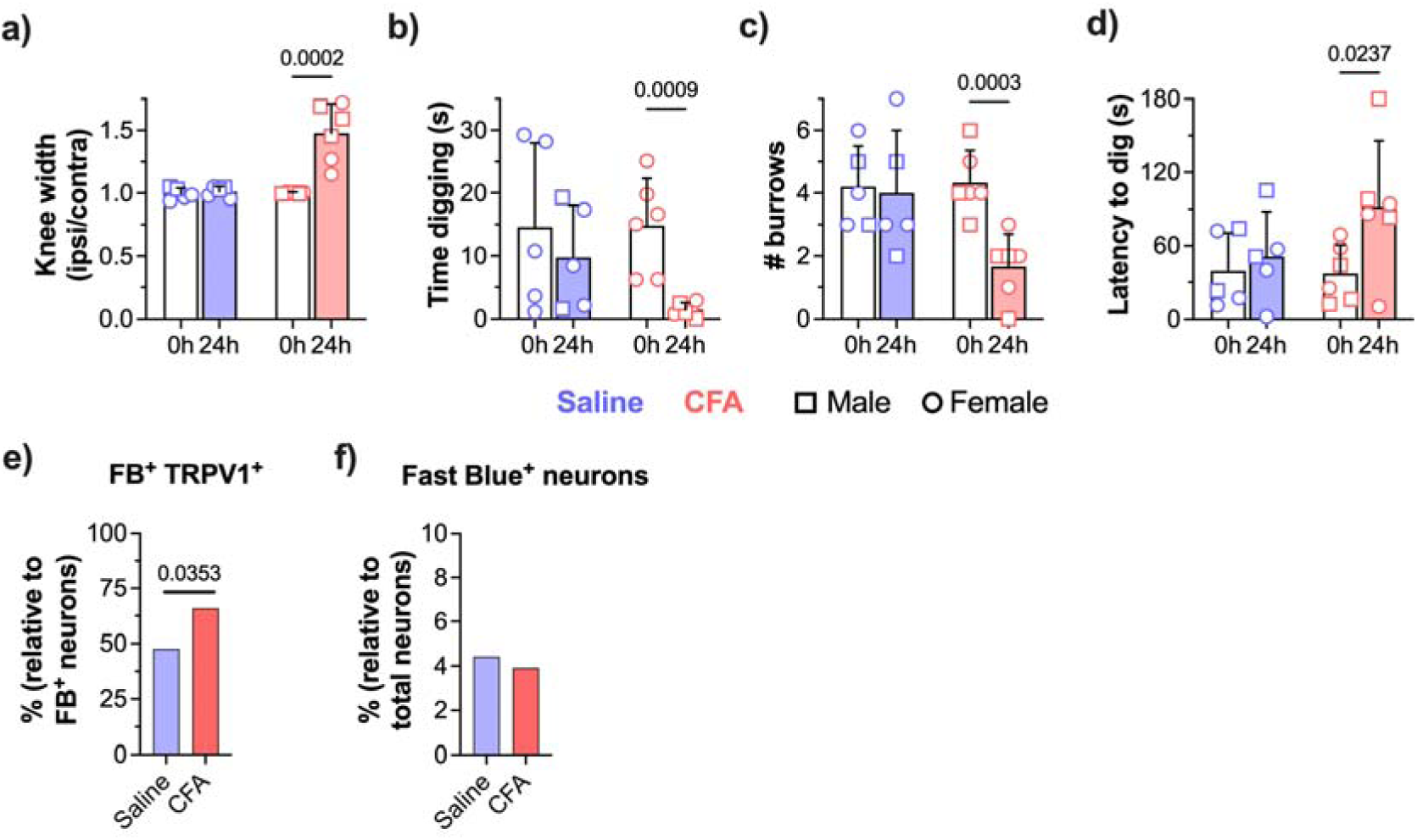
Decreased digging behaviour in mice undergoing CFA-induced knee inflammation and association with increased TRPV1 neuronal expression. **a)** Ratio of ipsilateral knee width to contralateral knee width on the day of injections and 24-hours after (saline, n = 5; CFA, n = 6). Two-way repeated measures ANOVA with Sidak’s multiple comparisons test; F (1, 9) = 18.2, P = 0.0021. **b)** Time spent digging before injections and 24 hours after (saline, n = 5; CFA, n = 6). Two-way repeated measures ANOVA with Sidak’s multiple comparisons test; F (1, 9) = 5.256, P = 0.0476. **c)** Number of visible burrows at the conclusion of the 3-minute digging test on the day of injections and 24 hours after (saline, n = 5; CFA, n = 6). Two-way repeated measures ANOVA with Sidak’s multiple comparisons test; F (1, 9) = 14.7, P = 0.0040. **d)** Latency of mice to dig on the day of injections and 24-hours after (saline, n = 5; CFA, n = 6). Two-way repeated measures ANOVA with Sidak’s multiple comparisons test; F (1, 9) = 6.589, P = 0.0303. **e)** Proportion of knee-innervating DRG neurons (L2-L5) that express TRPV1 (saline n = 19/42 neurons [FB], 951 neurons [total]; CFA n = 25/39 neurons [FB], 998 neurons [total]; 3 mice/group). Two-sided Fisher’s exact test. **f)** Proportion of knee-innervating (Fast Blue^+^) DRG neurons (L2-L5) (saline n = 42/951 neurons [total]; CFA n = 39/998 neurons [total]; 3 mice/group). Two-sided Fisher’s exact test. *P* values are shown in plots. In b, c and d, data shown as mean ± SD.

**Supplementary Figure 6.**
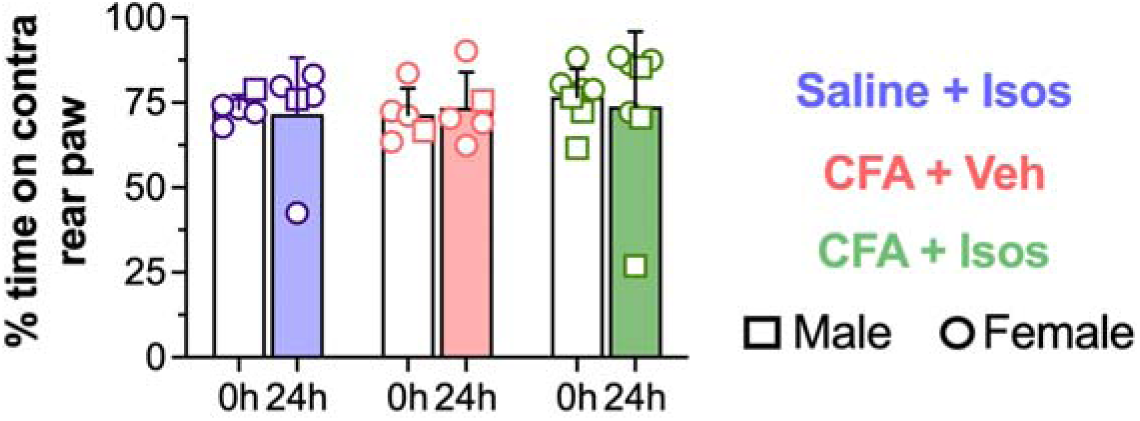
Time spent standing on the contralateral rear paw after treatment with isosakuranetin. Percentage of time spent standing on the contralateral rear paw at the conclusion of the 3-minute test on the day of injections and 24-hours after (saline+isos, n = 5; CFA+veh, n = 5; CFA+isos, n = 7). Two-way repeated measures ANOVA with Sidak’s multiple comparisons test; F (2, 14) = 0.1149, P = 0.8923. *P* values are not shown as statistical test did not show statistically significant differences between groups or time points. Data shown as mean ± SD.

## Notes

### Competing Interest Statement

The authors have declared no competing interest.

